# YlaN is an iron(II) binding protein that functions to relieve Fur-mediated repression of gene expression in *Staphylococcus aureus*

**DOI:** 10.1101/2023.10.03.560778

**Authors:** Jeffrey M. Boyd, Karla Esquilín-Lebrón, Courtney J. Campbell, Kylie Ryan Kaler, Javiera Norambuena, Mary E. Foley, Timothy G. Stephens, Gustavo Rios, Gautam Mereddy, Vincent Zheng, Hannah Bovermann, Jisun Kim, Arkadiusz W. Kulczyk, Jason H. Yang, Todd M. Greco, Ileana M. Cristea, Valerie J. Carabetta, William N. Beavers, Debashish Bhattacharya, Eric P. Skaar, Dane Parker, Ronan K. Carroll, Timothy L. Stemmler

## Abstract

Iron (Fe) is a trace nutrient required by nearly all organisms. As a result of the demand for Fe and the toxicity of non-chelated cytosolic ionic Fe, regulatory systems have evolved to tightly balance Fe acquisition and usage while limiting overload. In most bacteria, including the mammalian pathogen *Staphylococcus aureus*, the ferric uptake regulator (Fur) is the primary transcriptional regulator that controls the transcription of genes that code for Fe uptake and utilization proteins. YlaN was demonstrated to be essential in *Bacillus subtilis* unless excess Fe is added to the growth medium, suggesting a role in Fe homeostasis. Here, we demonstrate that YlaN is expendable in *S. aureus*; however, YlaN became essential upon Fe deprivation. A null *fur* allele bypassed the essentiality of YlaN. The transcriptional response of Fur derepression resulted in a reprogramming of metabolism to prioritize fermentative growth over respiratory growth. The absence of YlaN diminished the derepression of Fur-dependent transcription during Fe limitation. Bioinformatic analyses suggest that *ylaN* was recruited to Gram positive bacteria and once acquired was maintained in the genome as it co-evolved with Fur. Consistent with a role for YlaN in influencing Fur-dependent regulation, YlaN and Fur interacted *in vivo*. YlaN bound Fe(II) *in vitro* using oxygen or nitrogen ligands with an association constant that is consistent with a physiological role in Fe sensing and/or buffering. These findings have led to a model wherein YlaN is an Fe(II) binding protein that influences Fur-dependent regulation through direct interaction.

**Importance:** Iron (Fe) is an essential nutrient for nearly all organisms. If Fe homeostasis is not maintained, Fe can accumulate in the cytosol where it is toxic. Questions remain about how cells efficiently balance Fe uptake and usage to prevent imbalance. Iron uptake and proper metalation of proteins are essential processes in the mammalian bacterial pathogen *Staphylococcus aureus*. Understanding the gene products involved in Fe ion regulation, uptake, and usage, as well as the physiological adaptations that *S. aureus* uses to survive in Fe-depleted conditions, will provide insight into the role that Fe has in pathogenesis. These data will also provide insight into the selective pressures imparted by the mammalian host.

## Introduction

Iron (Fe) is an essential nutrient for nearly all organisms. Cytosolic Fe overload can cause toxicity through Fenton chemistry and/or the mismetalation of proteins (1). Because of this paradox, organisms must properly balance Fe uptake with Fe usage to ensure fitness and survival. *Staphylococcus aureus* requires Fe to proliferate and infect host tissues (2). The *S. aureus* genome codes for numerous Fe acquisition systems including the synthesis of two siderophores and one metallophore (3, 4). *In toto*, thus far, 2% of protein coding ORFs code for Fe uptake proteins (3, 5). Once acquired, Fe is used to build iron-sulfur clusters (FeS) and heme, as well as to metalate proteins. The process of building FeS clusters is essential, making Fe acquisition vital (6).

The ferric uptake regulator (Fur) is a master regulator of Fe uptake in bacteria (7). Few studies have examined the function of Fur in Gram-positive bacteria including phylum Firmicutes, in which *S. aureus* and *Bacillus subtilis* are members (8). In general, the accepted model for Fur regulation is that holo-Fur acts as a transcriptional repressor or activator (9). Fur can bind a variety of divalent metals *in vitro*, but it is thought that Fe(II) acts as the co-repressor *in vivo* (10, 11). Fur binds directly to operators and represses transcription of genes that code for proteins involved in Fe uptake during Fe replete conditions. When the bioavailable concentration of Fe is low, Fur is demetallated and repression is relieved. Fur often also controls the expression of a sRNA (*tsr25* in *S. aureus*) that helps to optimize Fe usage (12).

The *ylaN* gene is essential in *Bacillus subtilis* and the essentiality can be bypassed by growth with excess Fe salts, but the function of YlaN has not been described (13). Decreased *ylaN* expression results in phenotypes that mimic decreased expression of *sufCDSUB*, which codes the essential FeS cluster synthesis system. These findings led us to hypothesize that YlaN functions in Fe ion homeostasis. In this study, we created a Δ*ylaN* mutant in *S. aureus* and demonstrate that this mutation is lethal when cells are cultured in low Fe conditions. The growth defects of a Δ*ylaN* mutant were relieved by null mutations in *fur* or by fermentative growth. We demonstrate that YlaN binds to Fe(II) and that YlaN functions to relieve repression of Fur regulated genes during Fe limitation. Our findings support a model wherein YlaN is an Fe(II) binding protein that influences Fur-dependent regulation through a direct interaction.

## Results

### A *S. aureus* Δ*ylaN* mutant is defective in Fe ion processing

We created a Δ*ylaN::tetM* mutant in the community acquired methicillin-resistant *S. aureus* (CA-MRSA) isolate USA300_LAC (WT). The growth of the Δ*ylaN::tetM* strain was indistinguishable from that of WT in tryptic soy broth (TSB) medium (Figure S1). We also generated a *ylaN::Tn* strain in LAC that also grew similar to the WT in TSB. The metal ion chelators 2,2’-dipyridyl (DIP) and ethylenediamine-*N*,*N*’-bis(2-hydroxyphenylacetic acid) (EDDHA) have a high affinity for Fe (14, 15). DIP can cross the cell membrane, whereas the larger EDDHA likely does not (16, 17). Figure 1A displays the growth of the WT with the pCM28 (empty vector) or the Δ*ylaN::tetM* strain containing either pCM28 or pCM28_*ylaN* on tryptic soy agar (TSA) media with or without DIP or EDDHA. The Δ*ylaN::tetM* strain had a growth defect on both Fe-depleted media and the phenotype could be genetically complimented.

**Figure 1.**
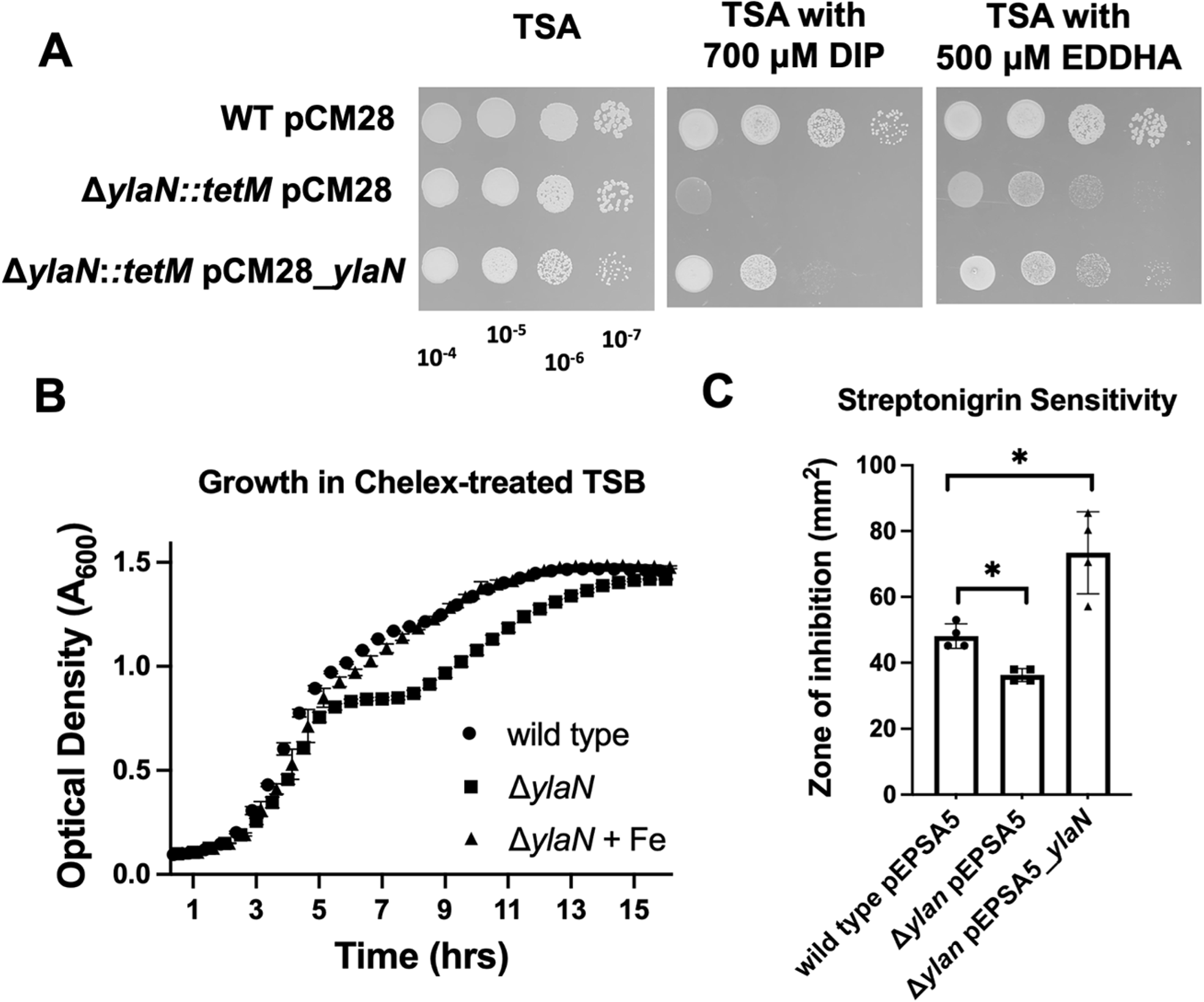
A role for YlaN in Fe ion homeostasis. **Panel A:** Growth of the WT (JMB1100) and Δ*ylaN::tetM* (JMB8689) with pCM28 or pCM28_*ylaN* on TSA-Cm with or without 2,2-dipyridyl (DIP) or EDDHA. Overnight cultures were grown in TSB, serial diluted, and spot plated. Plates were incubated for 18 hours before visualization. Pictures of a representative experiment are displayed. **Panel B:** Growth of the WT and Δ*ylaN::tetM* strains in Chelex-treated TSB medium supplemented with trace metals lacking Fe with and without 40 μM Fe(II). Data represent the average of biological triplicates and errors are displayed as standard deviations. **Panel C:** Streptonigrin sensitivity was monitored using TSB-Cm top-agar overlays enclosing the WT or Δ*ylaN::tetM* strains containing either pEPSA5 or pEPSA5_*ylaN*. The data in panels B and C represent the average of three biological replicates with standard deviations displayed. Student’s t-tests were performed on the data in panel C and * indicates p < 0.05.

To ensure that the growth defect was the result of decreased Fe ion availability and not decreased availability of an alternate metal ion, we treated tryptic soy broth (TSB) medium with Chelex-100 and then added back trace metals except Fe. The Δ*ylaN::tetM* strain had a growth defect in this medium and the growth defect was alleviated by the addition of Fe salts (Figure 1B). The Fe-dependent growth defect was apparent after approximately five hours of growth when glucose fermentation slowed and respiratory growth, which requires more Fe-dependent enzymes, was initiated (6).

The antibiotic streptonigrin, when combined with intracellular non-chelated Fe ions (sometimes referred to as “free Fe”) and electrons, causes double-stranded DNA breaks (18). Strains that are hypo- or hyper-active for Fe uptake are more resistant and more sensitive to streptonigrin, respectively (19). We created TSA overlays containing the WT with pEPSA5 (empty vector) or the Δ*ylaN::tetM* strain with pEPSA5 or pEPSA5_*ylaN*. The pEPSA5_*ylaN* vector placed *ylaN* under the transcriptional control of *xylRO* allowing for xylose induction of *ylaN* expression. We subsequently spotted streptonigrin and measured the zone of growth inhibition. The Δ*ylaN::tetM* strain was more resistant than the WT to growth in the presence of streptonigrin, suggesting that this strain had decreased streptonigrin associated Fe (Figure 1C). Over-production of *ylaN* made cells more sensitive to streptonigrin, suggesting that they had increased streptonigrin accessible Fe.

### Null mutations in *fur* suppress growth defect of the *ylaN* mutant in Fe limiting conditions

We conducted a suppressor screen to investigate YlaN essentiality under low Fe growth conditions. We individually plated ten independent cultures of the Δ*ylaN::tetM* strain on TSA medium containing 700 μM DIP. Colonies arose that contained second site mutations that permitted growth. We mapped these mutations, and nine strains contained a single nucleotide polymorphism (SNP) in the gene that codes for the ferric uptake regulator (Fur) resulting in the following variants: V29F (isolated twice), C103Y (isolated twice), E52X, T62M, R61I, R23H, and E11X (henceforth referred to as *fur*\*). The tenth strain had a guanine to cytosine transversion mutation in the 5’ untranslated region 13 base pairs upstream of the predicted translation start codon.

We linked a transposon (Tn) (SAUSA300_1452; *proC*) to the *fur\** mutation and then transduced the *fur*\* allele into the WT and Δ*ylaN::tetM* strains by selecting for the *proC::Tn*. As expected, some transductants had the *fur*\* allele while others retained the WT copy of *fur*. All isolates with *fur\** suppressed the growth defect of Δ*ylaN::tetM* on TSA medium with DIP, whereas those isolated with the WT *fur* allele did not (Figure 2A).

**Figure 2.**
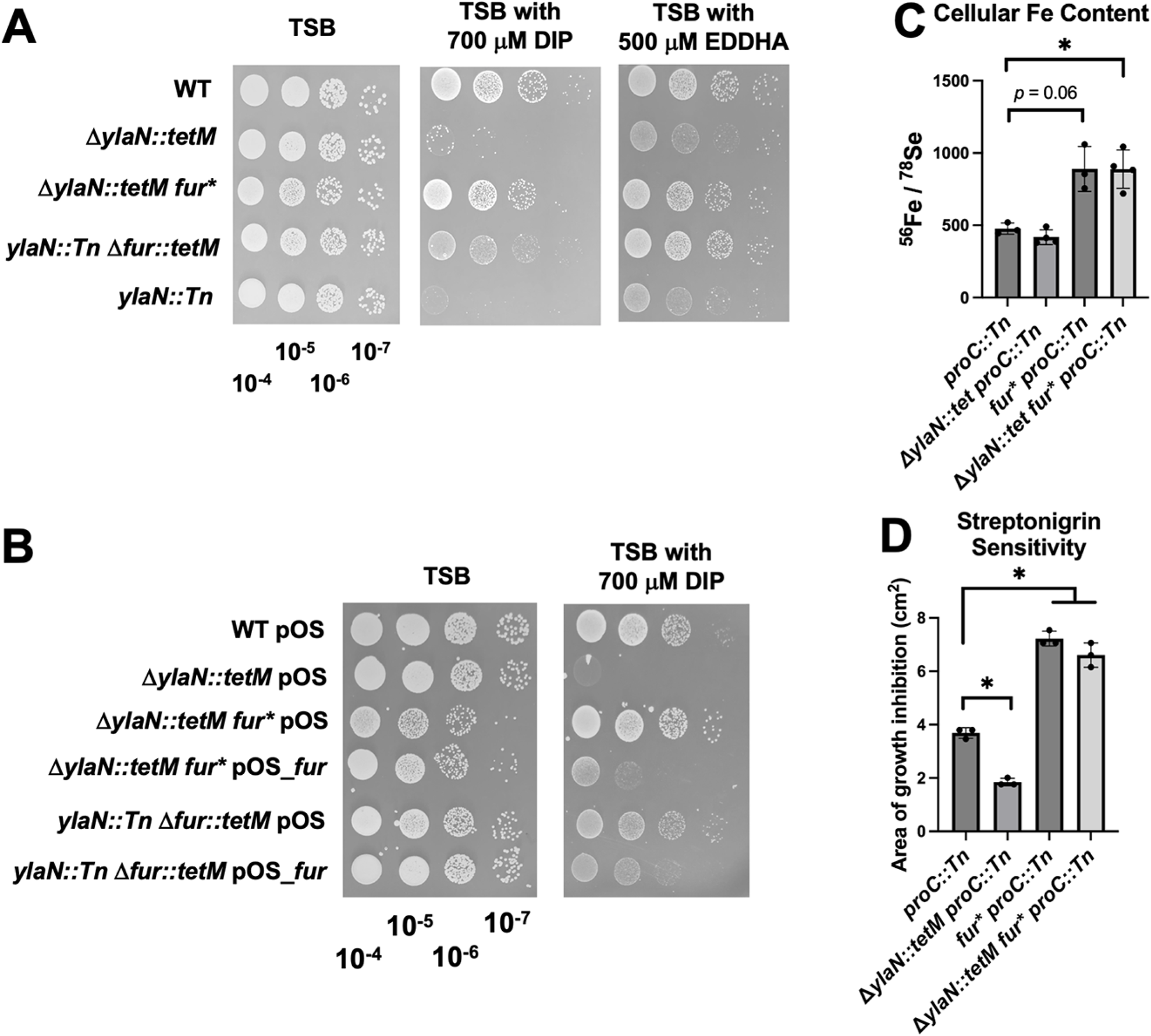
Null *fur* mutation bypasses the essentiality of YlaN in Fe-deplete conditions. **Panel A:** Overnight cultures of the WT (JMB1100), Δ*ylaN::tetM* (JMB8689), Δ*ylaN::tetM fur\** (JMB10678), *ylaN::Tn fur::tetM* (JMB10641), and *ylaN::Tn* (JMB8428) strains were serial diluted and spot plated on TSB media with or without 2,2-dipyridyl (DIP) or EDDHA. **Panel B:** Overnight cultures of the WT, Δ*ylaN::tetM*, Δ*ylaN::tetM fur\**, and *ylaN::Tn fur::tetM* strains containing either pOS or pOS_*fur* were serial diluted and spot plated on TSB-Cm media with or without DIP. **Panel C**: The ^57^Fe load was quantified in whole cells using ICPMS after culture in TSB medium. The ratio of ^57^Fe / ^78^Se is displayed for the *proC::Tn* (JMB10675), Δ*ylaN::tetM proC::Tn* (JMB10677), *fur* proC::Tn* (JMB10676), and Δ*ylaN::tetM fur* proC::Tn* (JMB10678) strains. **Panel D:** Streptonigrin sensitivity was monitored using top-agar tryptic soy agar overlays enclosing the *proC::Tn*, Δ*ylaN::tetM proC::Tn*, and *fur* proC::Tn*, and Δ*ylaN::tetM fur* proC::Tn* strains. Five μL of 2.5 mg mL^-1^ streptonigrin was spotted upon the overlays and the area of growth inhibition was measured after 18 hours of growth. Pictures of representative experiments are displayed in Panels A and B after 18 hours of growth. The data in panels C and D represent the average of three biological replicates with standard deviations displayed. Student’s t-tests were performed on the data in panels C and D. * indicates p < 0.05.

Two results are consistent with the hypothesis that the mutant *fur* alleles are null and recessive. First, we found that a Δ*fur::tetM* allele could suppress the DIP and EDDHA sensitivity phenotypes of a *ylaN::Tn* mutant (Figure 2A). Second, we were able to genetically complement the *fur\** and Δ*fur::tetM* alleles (Figure 2B).

We tested the hypothesis that the *fur\** allele was derepressing transcription of Fe uptake systems, and thereby increasing Fe import. We quantified the total Fe load in the WT and Δ*ylaN::tetM* strains after growth in TSB. As expected, there was a significant increase in Fe in the *fur*\* mutant strains (Figures 2C and S2). The *fur\** allele also increased streptonigrin sensitivity of the Δ*ylaN::tetM* mutant (Figure 2D). These data are consistent with the hypothesis that the *fur\** allele increased the total Fe load and the amount of intracellular non-chelated Fe.

We next tested the hypothesis that increased Fe uptake by introduction of the *fur\** allele suppressed the DIP sensitivity of the Δ*ylaN* strain. We introduced a *fhuC::Tn* mutation into the Δ*ylaN::tetM fur\** strain, which inactivated the siderophore-dependent Fe uptake systems (20, 21). The *fhuC::Tn* mutant had a growth defect on plates containing DIP as previously reported (Figure S3) (6). The *fur\** allele suppressed the growth defect of the Δ*ylaN::tetM fhuC::Tn* mutant suggesting that the null fur alleles were suppressing the growth defects of the Δ*ylaN* mutant by a mechanism other than promoting Fe uptake.

### Iron-deprivation or the loss of Fur alters the regulation of central metabolism

We developed and tested two non-mutually exclusive hypotheses to explain why a null *fur* allele suppressed the essentiality of YlaN under low Fe growth conditions: 1. The absence of functional Fur alters metabolism and thereby permits the Δ*ylaN* mutant to grow with a lower Fe allowance; and 2. YlaN functions to relieve Fur-dependent repression of transcription under low Fe growth conditions, which is bypassed by a null *fur* mutation.

We first tested the hypothesis that a null *fur* allele alters the regulation of metabolism to permit growth with a lower Fe allowance. We examined the effect of Fur and divalent metal starvation on the transcriptome. RNAs were quantified using RNA-sequencing after the WT and Δ*fur::tetM* strains were cultured in TSB with and without 120 μM DIP. This concentration of DIP was chosen because it does not cause a substantial defect in growth in either strain under the growth conditions utilized. Principal component analysis (PCA) of the regulons indicated that each strain and growth condition had a unique regulon and the individual regulons clustered (Figure S4). We verified the altered RNA abundances using quantitative PCR (qPCR) (Figure S5 and Table S1).

As expected, there was overlap between the WT vs. Δ*fur::tetM* and WT vs. WT + DIP regulons (Table S2). These regulons shared ≈ 30% of the genes with a log_2_ fold-change in abundance > ± 2 (Figure S6). The cause of the differences between these regulons is likely twofold. First, the addition of DIP competes for divalent metals altering the activity of divalent metal-dependent transcriptional regulators such as MntR and Zur. In support of this idea, there was increased transcription of divalent metal uptake genes (e.g. *abcA*, and *mntABC*) in the DIP regulon that were absent in the Fur regulon (Figure S7). Second, we did not add enough DIP to fully derepress Fur. In support of this, the transcription of Fe uptake genes (e.g. *iruO*, and *sstB*) was not fully derepressed in DIP treated cells (Figure S7).

Detailed analyses of the regulons showed that several genes coding nutrient uptake and energy generation enzymes had altered regulation in the Δ*fur::tetM* strain and DIP challenged WT (Figure S8). RNAs encoding Fe-uptake (*sbnB*) and Fe-requiring proteins (ribonucleotide reductase: *nrdD*) were increased and decreased, respectively, in the Δ*fur::tetM* strain and in the WT upon DIP challenge (Figures 3A and B and Table S2). There was decreased abundances of RNAs that code respiratory complexes (cytochrome oxidase: *qoxD*) and an increase in RNAs coding fermentation enzymes (formate dehydrogenase: *fdh*) in the Δ*fur::tetM* strain and in DIP challenged WT (Figures 3A and B and Table S2).

**Figure 3.**
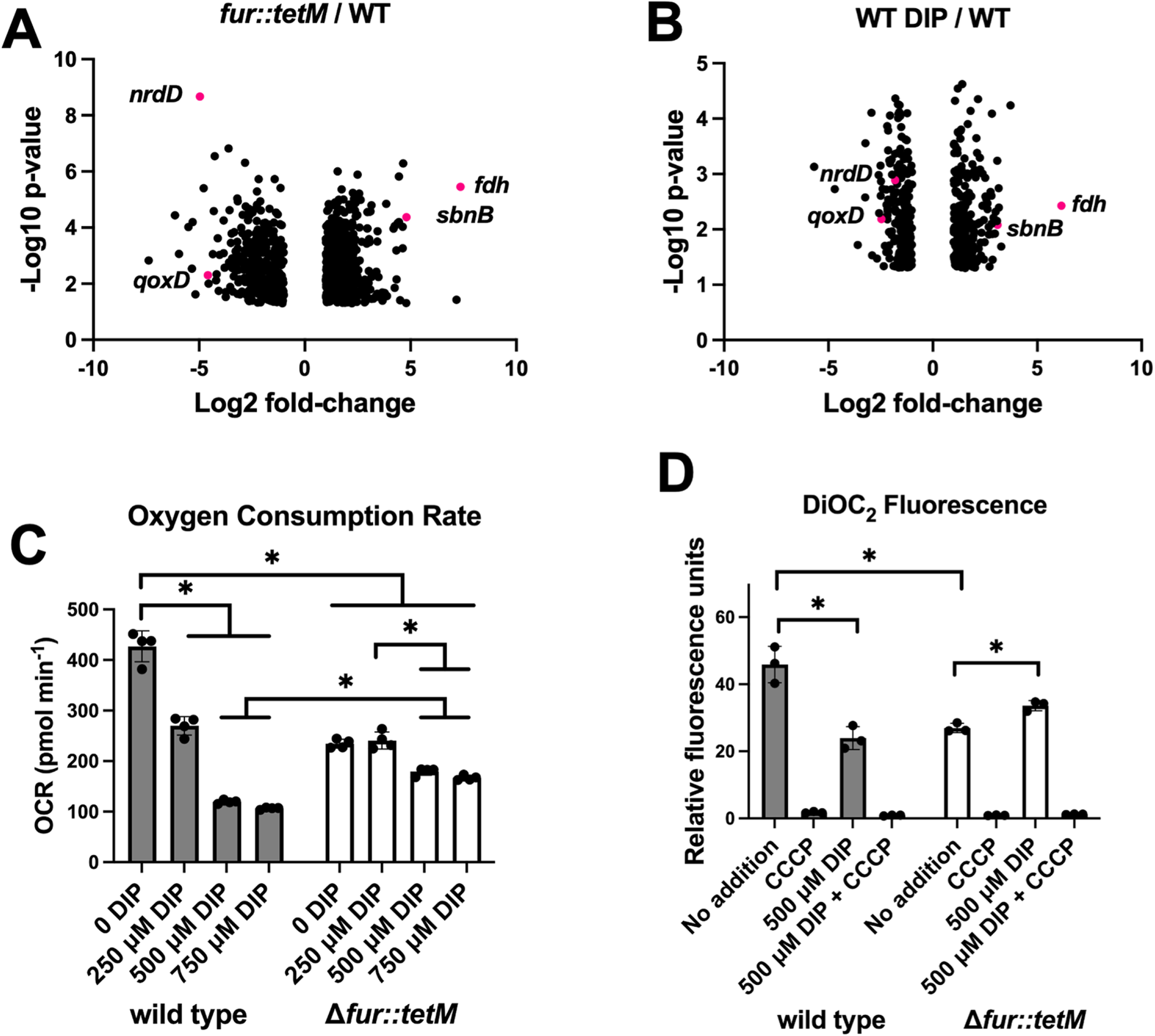
Transcriptome analyses suggest Fur-dependent transcriptional changes redirecting central metabolism upon Fe limitation. **Panel A:** Volcano plots of genes that have a Log2 fold change of ≥ 2 or ≤ −2 in the Δ*fur::tetM* (JMB1432) strain compared to the WT (JMB1100). **Panel B:** Volcano plots of genes that have a Log2 fold change of ≥ 2 or ≤ −2 in the 2,2-120 μM dipyridyl (DIP) challenged WT strain compared to unchallenged. **Panel C:** Dioxygen consumption rates (OCR) of the WT (JMB1100) and Δ*fur::tetM* (JMB1432) strains cultured in TSB media with and without 2,2-dipyridyl (DIP). OCR was monitored using a Seahorse Analyzer. **Panel D:** The membrane potentials of the WT and Δ*fur::tetM* strains were measured using the fluorescent dye 3′3′-diethyloxacarbocyanine iodide (DiOC_2_) after culture in TSB media with and without 2,2-dipyridyl (DIP). The data presented represent the average of biological triplicates. Standard deviations are displayed in Panels C and D and Student’s t-tests were performed and * indicates p < 0.05.

### Iron-deprivation or the absence of Fur decreases dioxygen respiration

To test the hypothesis that Fur downregulates cellular respiration in Fe-limiting conditions we quantified O_2_ consumption. First, we validated the technique by monitoring O_2_ consumption in a *hemB::Tn* mutant which cannot maturate cytochrome oxidases (22). The *hemB::Tn* strain had no detectable O_2_ consumption verifying that respiration is the primary consumer of O_2_ under the growth conditions examined (Figure S9A). We next examined the rates of O_2_ consumption using the WT and Δ*fur::tetM* strains cultured with varying concentration of DIP. The rate of O_2_ consumption decreased in the WT as a function of increasing DIP (Figure 3C). The basal rate of O_2_ consumption was decreased in the Δ*fur::tetM* mutant when compared to the WT. Dioxygen consumption was largely unaffected in the Δ*fur::tetM* strain when challenged with DIP consistent with the hypothesis that Fur derepression was responsible for the decreased O_2_ consumption in the DIP-challenged WT.

We further tested the ability of the WT and Δ*fur::tetM* strains to generate a membrane potential using the fluorescent dye 3′3′-diethyloxacarbocyanine iodide (DiOC_2_). When bacteria generate a membrane potential, DiOC_2_ accumulates intracellularly, resulting in a shift from emitting green fluorescence to emitting red fluorescence (23, 24). We validated the technique by analyzing the ability of DiOC_2_ to detect altered membrane potential between the WT and *hemB::Tn* strains. Whereas there was a red shift in fluorescence in the WT strain, very little red fluorescence was noted in the *hemB::Tn* mutant, indicative of a reduced membrane potential (Figure S9B). The addition of carbonyl cyanide m-chlorophenyl hydrazone (CCCP), which dissipates the transmembrane electric potential (Δψ) and ΔpH, resulted in decreased fluorescence in the WT (Figure S9B). We next examined the membrane potential in the WT and Δ*fur::tetM* strains after culture with and without 500 μM DIP. The membrane potential of the WT strain decreased upon co-culture with DIP (Figure 3D). The membrane potential was lower in the Δ*fur::tetM* strain compared to the WT. There was little change in the membrane potential in the Δ*fur::tetM* strain when co-cultured with DIP consistent with the hypothesis that Fur derepression was responsible for the decreased membrane potential in the WT.

### Iron-deprivation or the loss of Fur globally alters cellular metabolism

We performed genome-scale metabolic modeling using our RNA-sequencing data and O_2_ consumption measurements as modeling constraints (Table S3) (25). Consistent with these data, modeling analyses predicted decreased NADH dehydrogenase, succinate dehydrogenase, and cytochrome oxidase activity (Figure 4A), as well as decreased TCA cycle activity (Figure 4B) in iron limiting conditions. To validate these predictions, we experimentally measured the activity aconitase (AcnA), a TCA cycle FeS cluster requiring enzyme, in the WT Δ*fur::tetM* strains co-cultured with DIP (19). AcnA activity was decreased in Fe limiting conditions and nearly undetectable in the Δ*fur::tetM* strain (Figure 4C).

**Figure 4.**
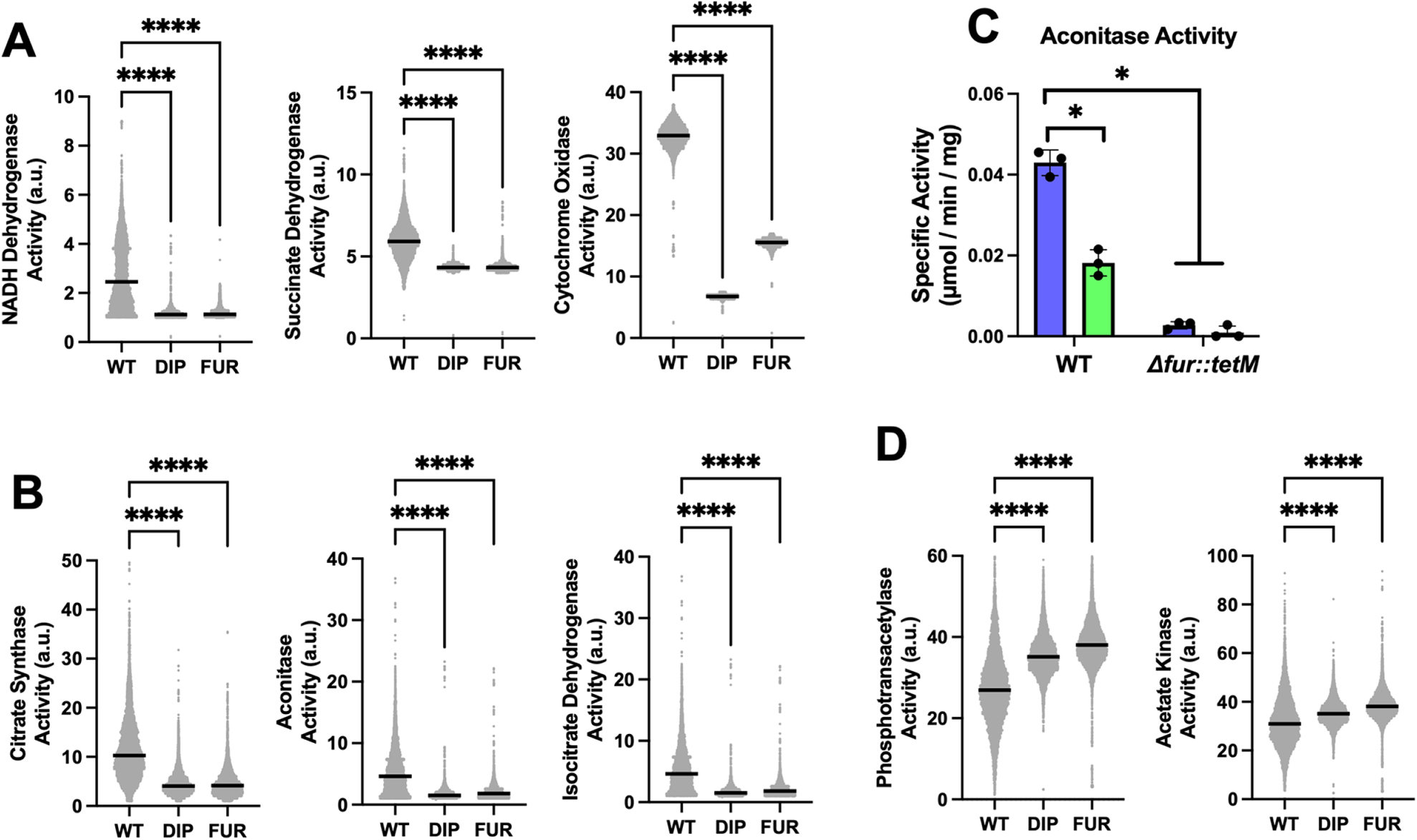
Genome-scale metabolic modeling analyses suggest Fe limitation induces fermentation. **Panels A and B:** Modeling analyses predict decreased respiratory activity (Panel A) and TCA cycle activity (Panel B). Flux balance analysis was performed on the iYS854 genome-scale metabolic model of *S. aureus* using transcriptome and dioxygen consumption measurements for WT, DIP-treated cells, and Δ*fur::tetM* strains as modeling constraints. **Panel C:** AcnA activity in cell-free lysates generated from the WT and Δ*fur::tetM* strains after four hours of culture with (green bars) and without (blue bars) 120 μM DIP. **Panel D:** Modeling analyses predict increased activity in acetate fermentation. For modeling analyses, data represent distributions for metabolic reaction activity simulations using 10,000 sampling points. Black lines represent median values for each distribution. For aconitase experiments, data represent the average of three biological replicates with standard deviations displayed. Kruskal-Wallis tests were performed on data in panels A, B, and C. Student’s t-tests were performed on the data in panel C. * indicates p < 0.05 and **** indicates p < 0.0001.

Modeling analyses further predicted several other global metabolic changes, including increased activity through acetate fermentation (Figure 4D), branched-chain amino acid biosynthesis, nucleotide biosynthesis, and the urea cycle. Modeling analyses also predicted decreased activity in lipid and cell wall biosynthesis and tRNA synthetase pathways (Table S3). Collectively, these data and modeling analyses are consistent with the hypothesis that upon Fe deprivation, Fur-dependent transcriptional repression is relieved, resulting in carbon flux being redirected to decrease reliance upon Fe-dependent enzymes, including respiratory complexes and TCA cycle enzymes, and increasing carbon flux through fermentation pathways to maintain redox homeostasis.

### Iron-deprivation or the absence of Fur alters metabolite abundances

To validate some of these modeling predictions, we cultured the WT and Δ*fur::tetM* strains, as we did for the transcriptomic experiments, before isolating metabolites and conducting untargeted metabolomic analyses. We also cultured the strains in the absence of O_2_ to force fermentation. The DIP treated WT and the Δ*fur::tetM* strain shared several metabolites that had altered abundances. For the aerobic samples, the Δ*fur::tetM* strain had more significantly altered metabolites than the DIP challenged WT (Tables 1 and S4), whereas the number of significantly altered metabolites were similar in the fermenting strains (Tables 2 and S4).

**Table 1.**
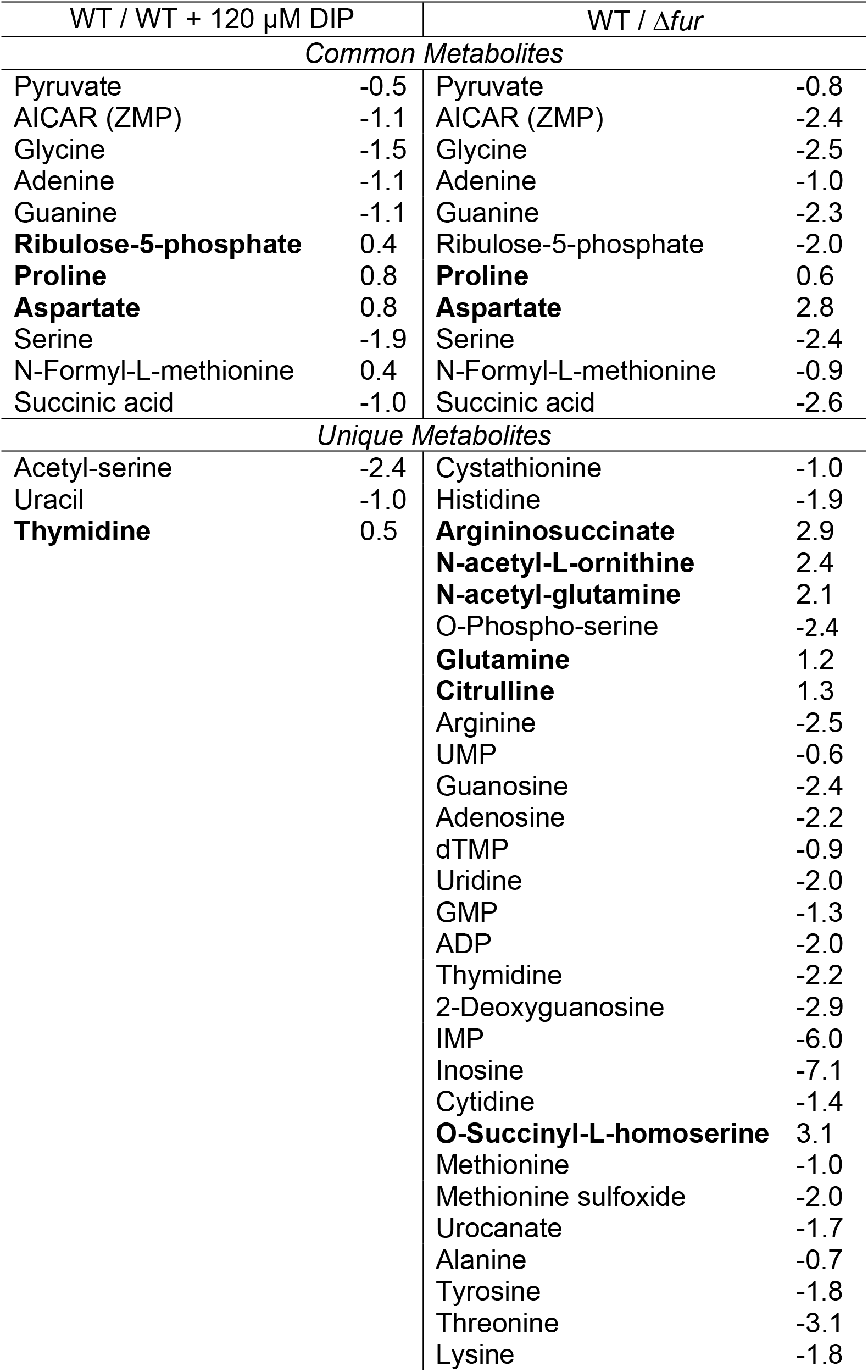
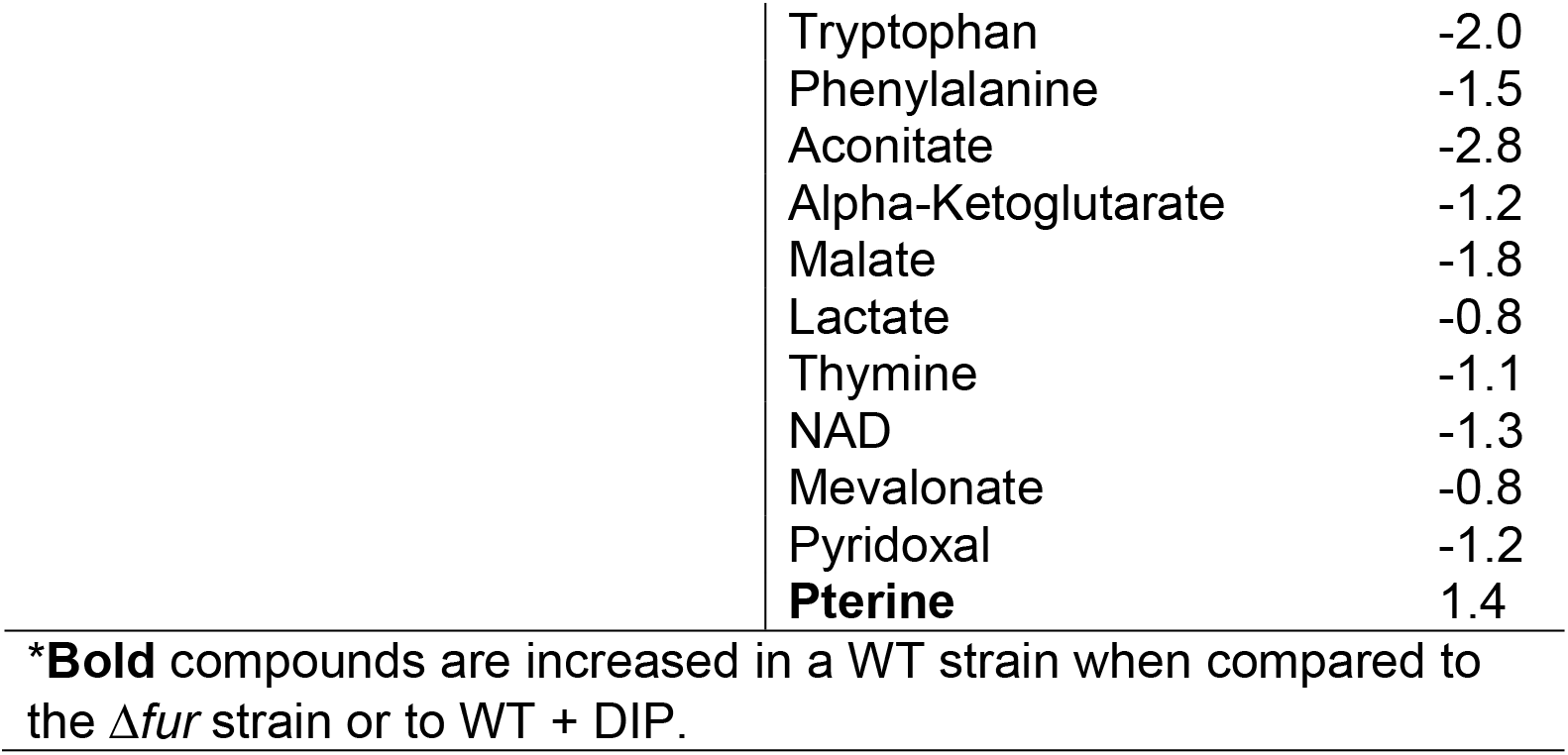
Log2 fold-changes of metabolites after aerobic growth*.

| Table 1. Log2 fold-changes of metabolites after aerobic growth* |  |  |  |
| --- | --- | --- | --- |
| WT / WT + 120 μM DIP | | WT / $\Delta fur$ | |
| Common Metabolites |  |  |  |
| Pyruvate | -0.5 | Pyruvate | -0.8 |
| AICAR (ZMP) | -1.1 | AICAR (ZMP) | -2.4 |
| Glycine | -1.5 | Glycine | -2.5 |
| Adenine | -1.1 | Adenine | -1.0 |
| Guanine | -1.1 | Guanine | -2.3 |
| **Ribulose-5-phosphate** | 0.4 | Ribulose-5-phosphate | -2.0 |
| **Proline** | 0.8 | **Proline** | 0.6 |
| **Aspartate** | 0.8 | **Aspartate** | 2.8 |
| Serine | -1.9 | Serine | -2.4 |
| N-Formyl-L-methionine | 0.4 | N-Formyl-L-methionine | -0.9 |
| Succinic acid | -1.0 | Succinic acid | -2.6 |
| Unique Metabolites |  |  |  |
| Acetyl-serine | -2.4 | Cystathionine | -1.0 |
| Uracil | -1.0 | Histidine | -1.9 |
| **Thymidine** | 0.5 | **Argininosuccinate** | 2.9 |
|  |  | **N-acetyl-L-ornithine** | 2.4 |
|  |  | **N-acetyl-glutamine** | 2.1 |
|  |  | O-Phospho-serine | -2.4 |
|  |  | **Glutamine** | 1.2 |
|  |  | **Citrulline** | 1.3 |
|  |  | Arginine | -2.5 |
|  |  | UMP | -0.6 |
|  |  | Guanosine | -2.4 |
|  |  | Adenosine | -2.2 |
|  |  | dTMP | -0.9 |
|  |  | Uridine | -2.0 |
|  |  | GMP | -1.3 |
|  |  | ADP | -2.0 |
|  |  | Thymidine | -2.2 |
|  |  | 2-Deoxyguanosine | -2.9 |
|  |  | IMP | -6.0 |
|  |  | Inosine | -7.1 |
|  |  | Cytidine | -1.4 |
|  |  | **O-Succinyl-L-homoserine** | 3.1 |
|  |  | Methionine | -1.0 |
|  |  | Methionine sulfoxide | -2.0 |
|  |  | Urocanate | -1.7 |
|  |  | Alanine | -0.7 |
|  |  | Tyrosine | -1.8 |
|  |  | Threonine | -3.1 |
|  |  | Lysine | -1.8 |

|  |  |
| --- | --- |
| Tryptophan | -2.0 |
| Phenylalanine | -1.5 |
| Aconitate | -2.8 |
| Alpha-Ketoglutarate | -1.2 |
| Malate | -1.8 |
| Lactate | -0.8 |
| Thymine | -1.1 |
| NAD | -1.3 |
| Mevalonate | -0.8 |
| Pyridoxal | -1.2 |
| **Pterine** | **1.4** |

\***Bold** compounds are increased in a WT strain when compared to the $\Delta fur$ strain or to WT + DIP.

**Table 2.** Log2 fold-changes of metabolites after anaerobic growth.

| WT / WT + 120 μM DIP |  | WT / Δ*fur* |  |
| --- | --- | --- | --- |
| Common Metabolites |  |  |  |
| Proline | 0.7 | Proline | 1.8 |
| Urocanate | 1.5 | Urocanate | 1.5 |
| Ornithine | 1.0 | Ornithine | 1.1 |
| Histidine | 2.3 | Histidine | 1.6 |
| Dihydroorotic acid | 2.4 | Dihydroorotic acid | -2.5 |
| 5-Aminolevulinate | 1.1 | 5-Aminolevulinate | 1.4 |
| Unique Metabolites |  |  |  |
| Orotate | 0.7 | N-acetyl-L-ornithine | 0.4 |
| N-carbamoyl-aspartate | 1.0 | O-Phospho-serine | -2.8 |
| Asparagine | 0.7 | Citrulline | 2.6 |
| Arginine | 0.4 | Glutamine | 0.7 |
| Pyruvate | -0.4 | Argininosuccinate | 3.3 |
| Phosphoenolpyruvate | 1.2 | N-acetyl-glutamine | 1.0 |
| Glyceraldehyde-3-phosphate | -2.2 | 2-Keto-3-deoxy-D-gluconate | 1.3 |
| Dihydroxyacetone phosphate | -2.4 | Diaminopimelic acid | -2.5 |
| Deoxycytidine | -0.7 | Thiamine | -0.5 |
| 2-Deoxyguanosine | -2.6 | dUMP | -2.4 |
| dUMP | -2.0 | CMP | 0.8 |
| AMP | 0.5 | AICAR (ZMP) | -1.1 |
| Pterine | 1.2 |  |  |
| NADH | 0.9 |  |  |
| Lysine | 0.6 |  |  |
| Acetyl-serine | -2.2 |  |  |
| Cystine | 1.0 |  |  |

\***Bold** compounds are increased in a WT strain when compared to the Δfur strain or to WT + DIP.

Independent of the presence of O_2_, the absence of Fur or the presence of DIP decreased pools of amino acids with nitrogen containing side chains (His, Gln, Pro) and decreased abundances of metabolites associated with nitrogen homeostasis (citrulline, ornithine, argininosuccinate, glutamine, N-acetyl-glutamine). Consistent with model predictions that nucleotide metabolism is induced by iron limitation (Table S3), aerobic co-culture with DIP, or Fur deletion, increased the abundance of adenine, guanine, thymine, and uracil. We hypothesize that this metabolic shift occurs to increase the production of the nitrogen containing metallophore, staphylopine, and the siderophores, staphyloferrin A and B. Consistent with this hypothesis, there were altered abundances of diaminopimelic acid, pyruvate, O-phospho-serine, serine, ornithine and histidine which are metabolic precursors used for the synthesis of these metallophores (5, 23, 26).

During aerobic growth, the Δ*fur::tetM* strain showed an increased accumulation of pyruvate, malate, α-ketoglutarate, and succinate. These metabolites were not altered in the fermenting cells suggesting that flux though the TCA cycle was altered during respiratory growth during Fe limitation.

### Fermentative growth bypasses the essentiality of YlaN during Fe deprivation

We tested the hypothesis that a null *fur* mutation alters metabolism and thereby bypasses the essentiality of YlaN in Fe-depleted media. We cultured the parent (*proC::Tn*), and isogenic *fur\**, Δ*ylaN::tetM,* and Δ*ylaN::tetM fur\** strains on solid or in liquid media with or without metal chelator in the presence and absence of O_2_. Since no other respiratory substrate was added to the media, anaerobic growth forces fermentation. The Δ*ylaN::tetM* strain displayed a growth defect with chelator when cultured aerobically (Figures 5A and C) but not when cultured anaerobically (Figures 5B and D). All four strains grew equally well with chelator when incubated anaerobically. These data are consistent with the hypothesis that the essentiality of YlaN in Fe deplete conditions can be bypassed by fermentative growth.

**Figure 5.**
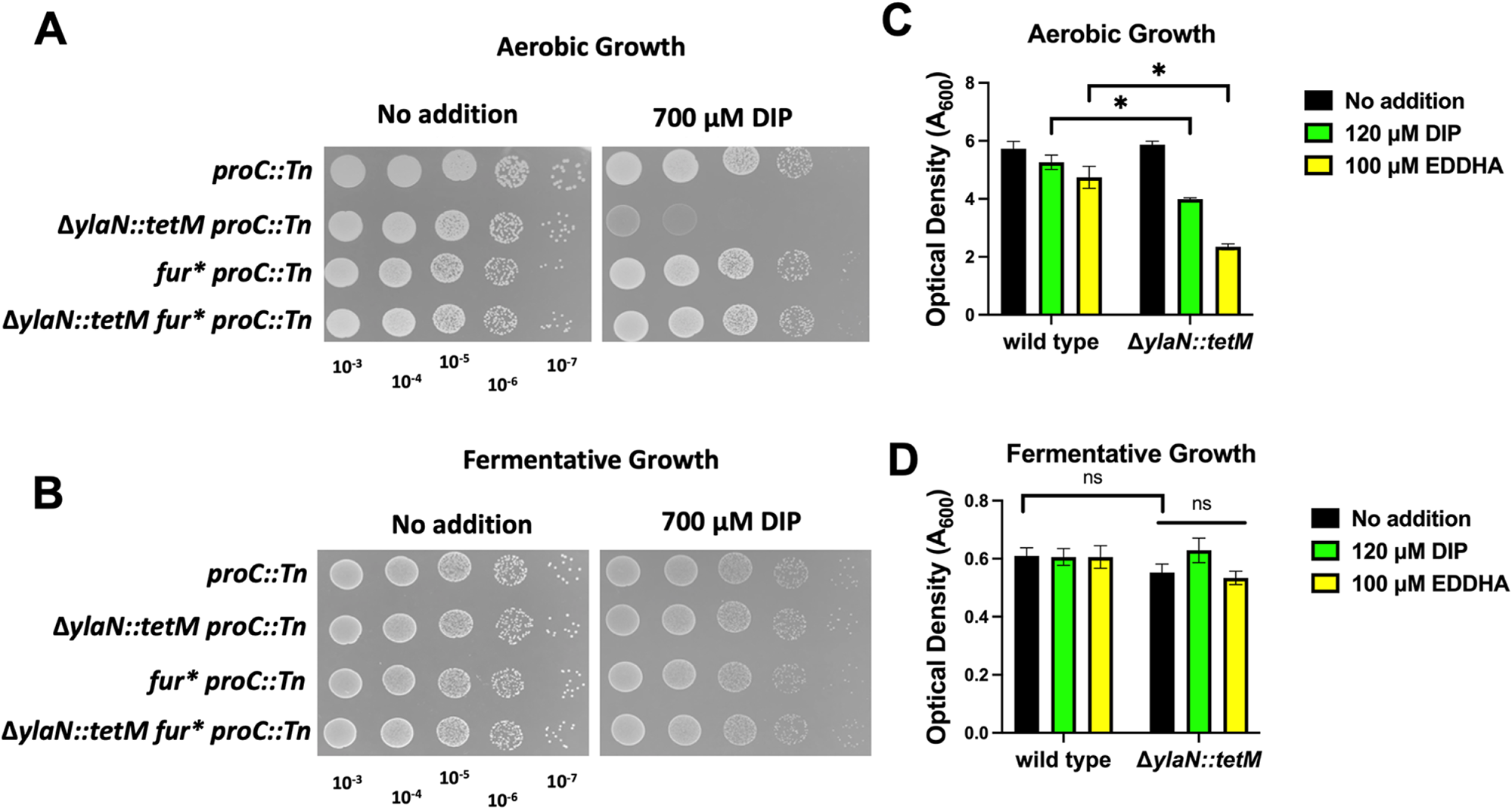
Fermentative growth bypasses the need for YlaN in Fe-deplete conditions. **Panels A and B:** The *proC::Tn* (JMB10675), Δ*ylaN::tetM proC::Tn* (JMB10677), *fur* proC::Tn* (JMB10676), and Δ*ylaN::tetM fur* proC::Tn* (JMB10678) strains were cultured overnight before serial dilution and spot plating on TSA with and without 2,2-dipyridyl (DIP). The cultures displayed in panel A were incubated aerobically and the cultures in panel B were incubated anaerobically in the absence of a terminal electron acceptor to force fermentative growth. Photographs of a representative experiment are shown. **Panels C and D**: The WT (JMB1100) and Δ*fur::tetM* (JMB1432) strains were inoculated into TSB medium with and without DIP or EDDHA before culturing overnight aerobically or anaerobically in the absence of a terminal electron acceptor. Culture optical densities were recorded after 18 hours of growth. The data displayed represent the average of biological triplicates and standard deviations are displayed. Student’s t-tests were performed on the data and * indicates p < 0.05.

### YlaN is required to relieve Fur-dependent repression of transcription

We tested the second hypothesis that YlaN is required to relieve Fur-dependent repression of transcription under low Fe growth conditions. The *fhuC*, *isdC*, *tsr25*, and *sbnA* genes are regulated by Fur (Table S2 and Figure S10) and have Fur-boxes in their promoters suggesting direct Fur regulation. Consistent with this, *fhuC*, *isdC,* and *tsr25* transcriptional reporters were more active in the Δ*fur::tetM* strains. We were unable to transduce the *sbnA* reporter into the *fur* mutant. All four promoters had increased activity in the WT upon growth with DIP (Figures 6A-D). In contrast, none of the promoters responded to DIP in the Δ*ylaN::tetM* mutant.

**Figure 6.**
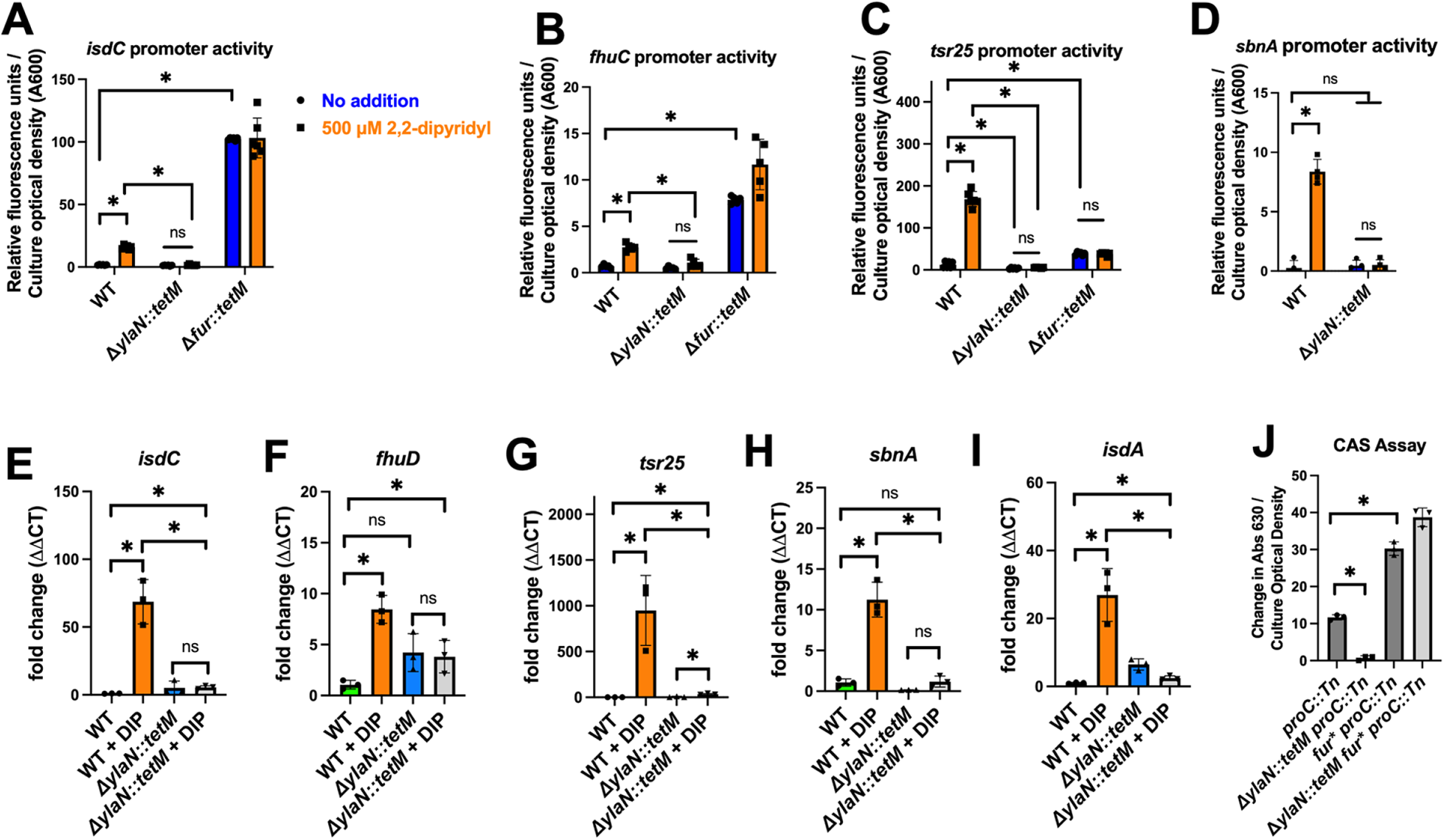
The presence of YlaN promotes Fur-dependent derepression in Fe-deplete conditions. **Panels A-D:** The transcriptional activities of the *isdC*, *fhuC, tsr25*, and *sbnA* promoters were measured in the WT (JMB1100), Δ*ylaN::tetM* (JMB8689), and Δ*fur::tetM* (JMB1432) strains after culture in TSB-Cm media with (blue bars) and without (orange bars) 500 µM 2,2-dipyridyl (DIP). We were unable to mobilize the sbnA transcriptional reporter into the *fur::tetM* strain. The data displayed represent the average of biological quintets and standard deviations are displayed. **Panels E-I:** Abundances of RNAs expressed from select Fur-regulated genes in the WT and *ylaN::tetM* strains were determined using quantitative PCR after culture in TSB media with and without 500 µM DIP. The data displayed represent the average of biological triplicates and standard deviations are displayed. **Panel J:** The ability of cell free culture spent medium to compete with chrome azurol S (CAS) for Fe was measured spectrophotometrically. Cell free spent culture medium was harvested from the *proC::Tn* (JMB10675), Δ*ylaN::tetM proC::Tn* (JMB10677), *fur* proC::Tn* (JMB10676), and Δ*ylaN::tetM fur* proC::Tn* (JMB10678) strains after culture in Chelex-treated TSB. The data displayed represent the average of biological triplicates with standard deviations shown. Student’s t-tests were performed on the data and * indicates p < 0.05.

We next quantified RNA transcripts corresponding to *isdC*, *fhuD*, *tsr25*, *sbnA*, and *isdA* in the WT and Δ*ylaN::tetM* strains after growth with and without DIP. All transcripts increased in the WT strain upon DIP challenge; however, none of these transcripts significantly responded to DIP challenge in the Δ*ylaN::tetM* strain (Figures 6E-I).

We next monitored siderophore production using chrome azurol S (CAS). Fe-bound CAS is blue in color. The addition of siderophores competes with CAS for the Fe resulting in apo-CAS, which is orange. We cultured the parent (*proC::Tn*), Δ*ylaN::tetM*, *fur\**, and Δ*ylaN::tetM fur\** strains in TSB before measuring CAS competition using spent media. The spent media from the strains containing the *fur*\* mutations were better able to compete for Fe than the parent (Figure 6J). The spent media from the Δ*ylaN::tetM* strain did not alter the absorbance of the CAS-Fe complex. These data are consistent with the hypothesis that the presence of YlaN functions to relieve Fur-dependent repression of transcription upon Fe limitation, which can be bypassed by a null *fur* mutation.

### YlaN and Fur likely co-evolved

We compared the conservation of *ylaN* and *fur* in bacterial genomes. The presence of *ylaN* is restricted to Firmicutes, forming a clear monophyletic clade (Figure 7). The Fur sequences from YlaN-positive genomes are paraphyletic, with the sequences from YlaN-negative genomes having diverged early in this group. This suggests that YlaN was lost from these lineages during the early stages of its evolution. There are only six sequences in the tree (all Firmicutes) that are from YlaN-positive genomes but are not part of the monophyletic clade of Firmicutes Fur sequences. These sequences are positioned with non-Fur sequences in the tree (e.g., Zur and PerR) and are likely to be PerR. They were recruited into the tree during the DIAMOND (sequence similarity-based) search, and thus support the singular origin of YlaN in Firmicutes.

**Figure 7.**
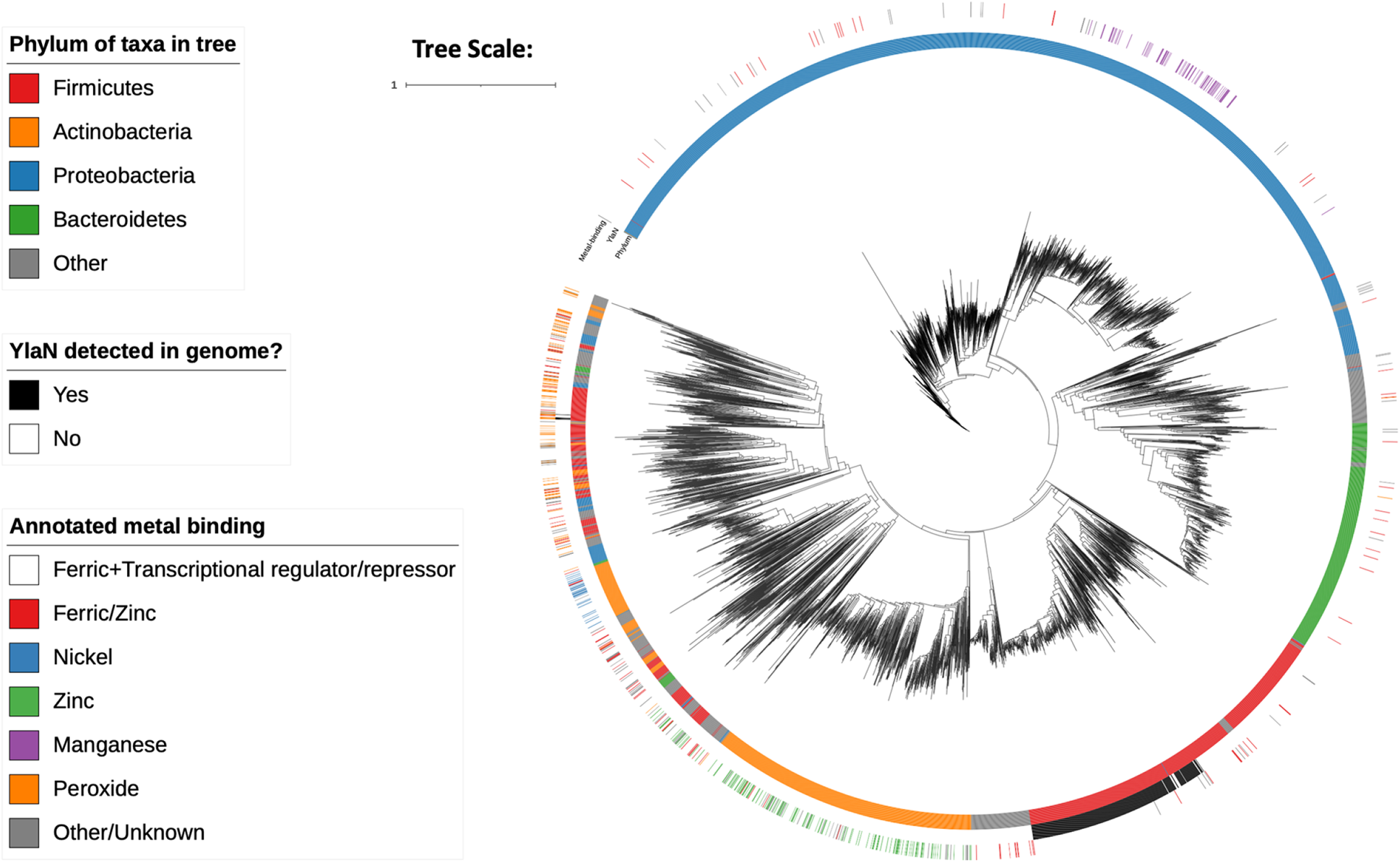
Phylogeny of representative bacterial Fur sequences. Tree (midpoint rooted) constructed using fastme from Fur sequences, down sampled to a genera level, extracted from high-quality representative or reference bacterial genomes. For each sequence in the tree, the colors in the surrounding rings represent the (inner ring) phylum, (middle ring) presence or absence of YlaN in the genome from which the Fur sequence was derived, and (outer ring) annotated metal-binding affinity. A legend showing the colors used in the surrounding rings is shown on the left side of the image, as is the branch length scale.

Comparison of the Firmicutes Fur and YlaN phylogenies (Figure S11 and S12) demonstrates that whereas there is significant discordance between both trees, generally, many of the closely related groups of sequences are present in both trees (represented by the colored lines) and the major differences arise from the early diverging nodes which are weakly supported (i.e., have bootstrap support <95%). This result suggests that Fur and YlaN have been co-evolving, although given the lack of phylogenetic signal for the early diverging nodes in both protein trees, this hypothesis is currently provisional.

### *Bacillus subtilis* YlaN and Fur and interact *in vivo*

The findings that the presence of YlaN alters the ability of Fur to respond to Fe limitation and that YlaN and Fur likely co-evolved in the Firmicutes led to the hypothesis that these two proteins physically interact with each other. We performed an immunoaffinity purification-mass spectrometry analysis of YlaN in non-pathogenic *B. subtilis*, which like *S. aureus*, is a member of the phylum Firmicutes. We chose to work in *B. subtilis* because the immunopurification protocol conditions are optimized (27) and for safety concerns since the sample preparation can create aerosols. We created a strain encoding an N-terminally fused YFP-YlaN from the *amyE* locus, which was under the transcriptional control of an IPTG-inducible promoter. YFP-YlaN was immunopurified from both logarithmic and stationary phase cells and co-isolated proteins were identified by mass spectrometry. Fur was the most abundant protein co-isolated with YFP-YlaN in both exponential and stationary phase (Table S5). The FeS cluster synthesis machinery (SufCDSUB), FeS cluster assembly factors (SufA and SufT), and Fe-requiring proteins (i.e., GlcF, NarH, QcrA, and CitB) were also identified as interacting partners. These data support the hypothesis that YlaN interacts with Fe utilizing proteins.

Using the AlphaFold2 structural models for *S. aureus* YlaN and Fur, we were able to model a potential interaction configuration (Figure S13). In these models, the highly conserved residues of YlaN helix two (EEVLDTQMFG) interact with the DNA binding domain of Fur.

### *Staphylococcus aureus* YlaN binds Fe(II) with a physiological relevant affinity

We tested the hypothesis that YlaN binds Fe(II) using a competition assay (28–30). This assay measures Fe(II) binding affinity of a protein under heterogenous solution conditions, which better mimics *in vivo* conditions. We measured changes in the UV-Vis signal at 365 nm from the apo-form of the chelator Mag-Fura-2, which decreases upon Fe(II) loading of the chelator. Binding analyses were completed on two independently prepared protein samples. Data were simulated to deconvolute the protein-metal binding relative to the known chelator metal binding affinity (for Mag-Fura-2, Fe(II) binding *K*_d_ = 6.5 ± 0.5 μM). The Fe(II) binding affinity for YlaN was best fit by one independent *K*_d_ at 2.07 ± 0.46 μM (Figure 8A).

**Figure 8.**
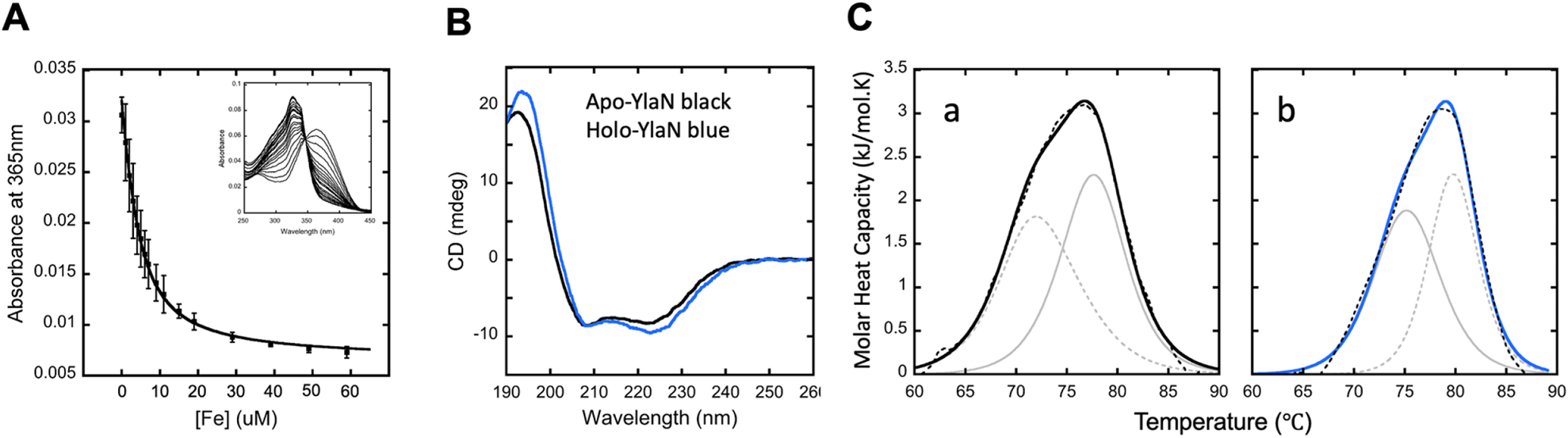
YlaN is an Fe(II) binding protein. **Panel A:** Fe binding affinity to YlaN measured using Mag-Fura-2 within a competition-based assay. Average titration spectral points of Fe(II) ions into YlaN with overall simulation in solid line. Mag-fura-2 to protein ratios were varied and 1:1 data is shown. Spectra were collected in duplicate using independent samples to ensure spectral reproducibility. **Panel B:** Circular dichroism (CD) Structural results for apo and Fe(II) loaded holo-YlaN. Representative CD spectra comparing apo- (black) and holo- (blue) YlaN. **Panel C:** Differential scanning calorimetric spectra for apo- and Fe(II) loaded holo-YlaN. Overall simulation of molar heat capacity vs. temperature spectra for apo- (a) and holo- (b) YlaN. Overall simulations are displayed in solid lines, with individual simulated features in gray and raw data shown in a black dashed line.

### Fe(II) binding to YlaN alters its secondary structure and thermal stability

We examined the impact of Fe(II) binding on the secondary structure of YlaN using circular dichroism (CD) spectroscopy. Comparison of average CD spectra of the apo- and holo-YlaN show some change between the two protein states (Figure 8B), with similar negative features at both 207 and 223 nm; wavelengths which are attributed to ⍺-helical structures (31). These negative features become more prominent upon Fe(II) binding. Simulation results are shown in Table S6. These results show an overall trend of increasing helical structure coupled with Fe(II) binding. The helical content of Fe(II) loaded protein is slightly increased (from 31% in the apo state to 40% in the holo), while the relative values of the additional structural elements only slightly decrease, suggesting that upon Fe(II) binding, YlaN increases its overall helical structure. Root-mean-square deviations (RMSD) between theoretical and empirical data for both apo- and holo-YlaN CD spectra are consistently low, suggesting simulations accurately predict the secondary structure change for each construct.

We examined the thermal stability of apo- and Fe(II) bound YlaN using differential scanning calorimetry (DSC). The apo-protein had a two-phase thermal stability profile (Figure 8C and Table S7). Melting temperatures for apo-YlaN were 72.0 and 77.5 °C. Fe(II) binding to YlaN increased the melting temperatures; the lower *T*_m_ was increased to 75.3 °C while the higher *T*_m_ value was increased to 79.8 °C, indicating that Fe(II) binding increased the protein’s thermal stability.

### The YlaN Fe(II) binding environment consists of only oxygen and/or nitrogen ligands

We probed the structure and electronic properties of Fe(II)-loaded YlaN to determine bound-metal coordination geometry. Structural details for Fe(II) bound to YlaN were measured using Fe *k*-edge X-Ray Absorption Spectroscopy (XAS) (32). Normalized X-ray Absorption Near Edge Spectroscopy (XANES) data was used to determine the average metal oxidation state, spin state, and ligand coordination symmetry (28). The XANES edge shape (Figure 9A, left) and first inflection edge energy at around 7123 eV (Table S8) suggest Fe-loaded YlaN contains only Fe(II). The pre-edge feature (Figure 9A, right) between 7110-7115 eV represents the Fe(II) 1s-3d electron transition. The area under the curve (Table S8) indicates protein bound Fe(II) is held in a highly symmetric 6-coordinate metal ligand environment.

**Figure 9.**
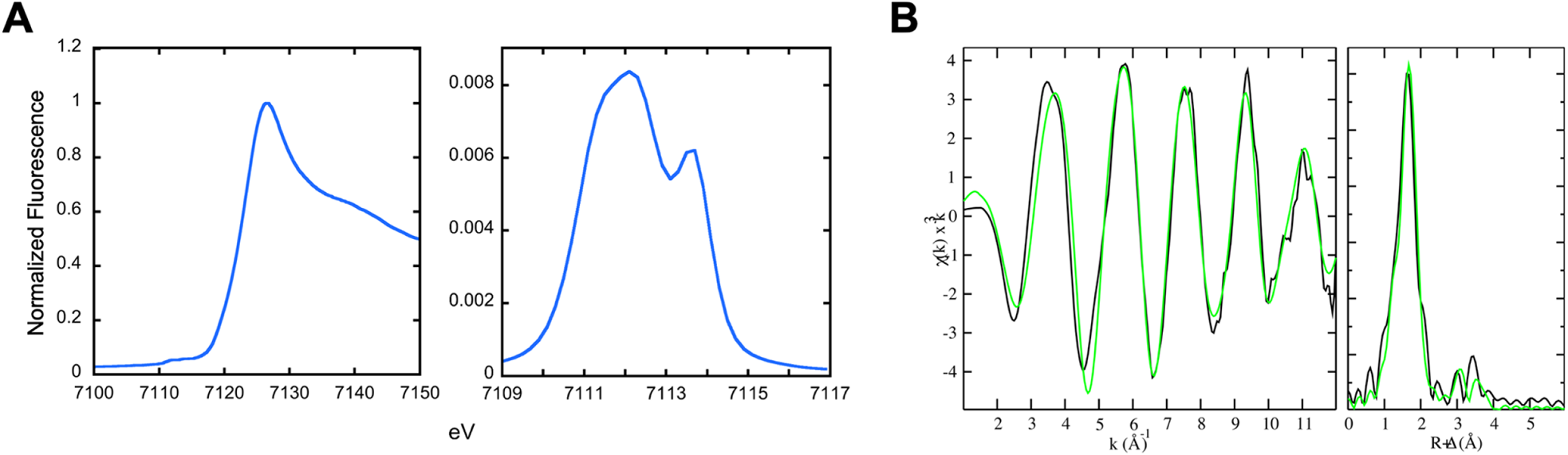
Examining the YlaN Fe(II) ligand environment. **Panel A:** Normalized XANES for Fe(II) bound to YlaN. Representative spectra for Fe XANES for Fe-YlaN (left) and extended display of Fe-XANES pre-edge region (right). **Panel B:** EXAFS analyses of Fe(II) bound YlaN. Raw EXAFS, Fourier transforms of EXAFS, and spectral simulations for Fe(II) bound YlaN. Fe EXAFS data and simulation of YlaN. Raw k^3^ weighted EXAFS data (left) and phase shifted Fourier transform (FT) (right) are shown. Raw data is shown in black, and simulations in green.

The strength of XAS is the high accuracy seen in the metal-ligand bond length values obtained by simulating the Extended X-Ray Absorption Fine Structure (EXAFS) portion of the XAS spectrum. In review, simulations of EXAFS data provide metal-ligand bond lengths at an accuracy of ± 0.02 Å, while information regarding metal-ligand coordination numbers are obtained at an accuracy of ± 1.0 and ligand identity narrowed to within one row of the periodic table (32). All Fe data were fit out to a *k*-space value of 12.5 Å^-1^ to eliminate high energy noise. Calibration from Fe(II) and Fe(III) theoretical model compounds were used for E_0_ and Sc parameters. E_0_ values for Fe-O, Fe-N, Fe-C were set at −10 eV and a scale factor of 0.95 was used to fit the data. Figure 9B shows the raw and simulated EXAFS and Fourier Transform (FT) of the EXAFS. Best fit simulations are listed in Table S9, which fit O/N as nearest neighbor ligands, followed by C as long range ligands. These data are consistent with a highly symmetric, octahedral Fe(II) coordination environment consisting of O/N ligands. The presence of C ligand scattering as a long-range interaction indicates Fe(II) in the sample with higher order, consistent with attachment to amino acids, as opposed to being bound adventitiously. Under these parameters, the average Debye Waller Factor and F’ values, measures of absorber-scatter bond disorder, and overall simulation convergence between theoretical and empirical data, were highly favorable.

## Discussion

We initiated this study after data produced by Peters *et al*. suggested that YlaN had a role in Fe ion homeostasis in *B. subtilis*. Unlike *B. subtilis*, YlaN is not essential in *S. aureus* under standard growth conditions. This difference in essentiality may be explained by the metabolic potential of these two organisms. Whereas *S. aureus* is a robust glucose fermenter in standard laboratory media, *B. subtilis* is inefficient at fermenting glucose in the absence of a respiration substrate (33, 34). In general, respiratory growth utilizes more FeS cluster proteins than fermentative growth (35). In support of this, *S. aureus* has decreased transcription of Fe uptake systems and imports less Fe when cultured in conditions that favor fermentative growth over respiratory growth (36). It is likely that *S. aureus* temporarily promotes a fermentative metabolism in low Fe conditions to prioritize the metalation of essential Fe enzymes, while sacrificing the Fe-demanding, but high ATP yielding respiratory growth.

We demonstrate that a null *fur* mutation bypasses the need for YlaN under Fe limiting conditions. The reason for this appears to be twofold. First, in the absence of YlaN, Fur does not properly derepress transcription during Fe limitation, which is bypassed by the introduction of a null *fur* allele. Second, in the absence of Fur, cells switch to a fermentative metabolism. Consistent with our findings herein, it was previously demonstrated that growth with DIP, or the absence of Fur, resulted in increased lactate fermentation (37). A second study found that growing with low Fe decreased abundances of metabolites associated with the TCA cycle and increased abundances of fermentation byproducts including acetate and lactate (36). We discovered that forcing fermentative growth by anaerobic incubation decreased the Fe-dependent growth defect of the Δ*ylaN* strain. These data are consistent with a model wherein YlaN functions to relieve Fur-dependent repression of transcription during Fe limitation. In the absence of YlaN, *S. aureus* cannot trigger the Fur-dependent switch to an Fe-restrictive fermentative metabolism resulting in a severe growth defect.

Bioinformatic analyses demonstrated that *ylaN* was recruited to a *fur*-containing genome and was then retained long-term, apparently shaping the evolutionary trajectory of *fur*. It is currently unknown if the *S. aureus* Fur binds Fe(II); however, YlaN does bind Fe(II) and the data here support the hypothesis that YlaN functions in Fur-dependent regulation. In support of this, YlaN and Fur interact *in vivo*. Modeling data predict that YlaN binds to Fur at the same location where it binds to DNA. The internal concentration for Fe(II) in *S. aureus* cells is unknown, but it is predicted to be *≈*6 μM in aerobically cultured *E. coli* (38). YlaN binds Fe(II) with an affinity (*K*_D_ *≈*2 μM) that is consistent with it being holo and apo in Fe replete and deplete conditions, respectively. The Δ*ylaN* mutant has less streptonigrin associated Fe suggesting that it is not functioning solely as an Fe buffer or Fe chaperone. The manuscript by Demann et al., which was co-submitted with this manuscript, demonstrates that the YlaN prevents the DNA binding activity of Fur (39). These data have resulted in a working model (Figure 10) wherein during Fe replete conditions, holo-Fur binds to operators of genes utilized in Fe uptake and represses transcription. During Fe limitation, YlaN interacts with Fur, resulting in derepression of Fur-regulated genes, as well as decreased expression of genes coding Fe requiring proteins. The latter is likely indirect regulation possibly requiring the *tsr25* sRNA. Ultimately, this results in the derepression of Fe uptake systems and a reprogramming of central metabolism to compete in a low Fe environment.

**Figure 10.**
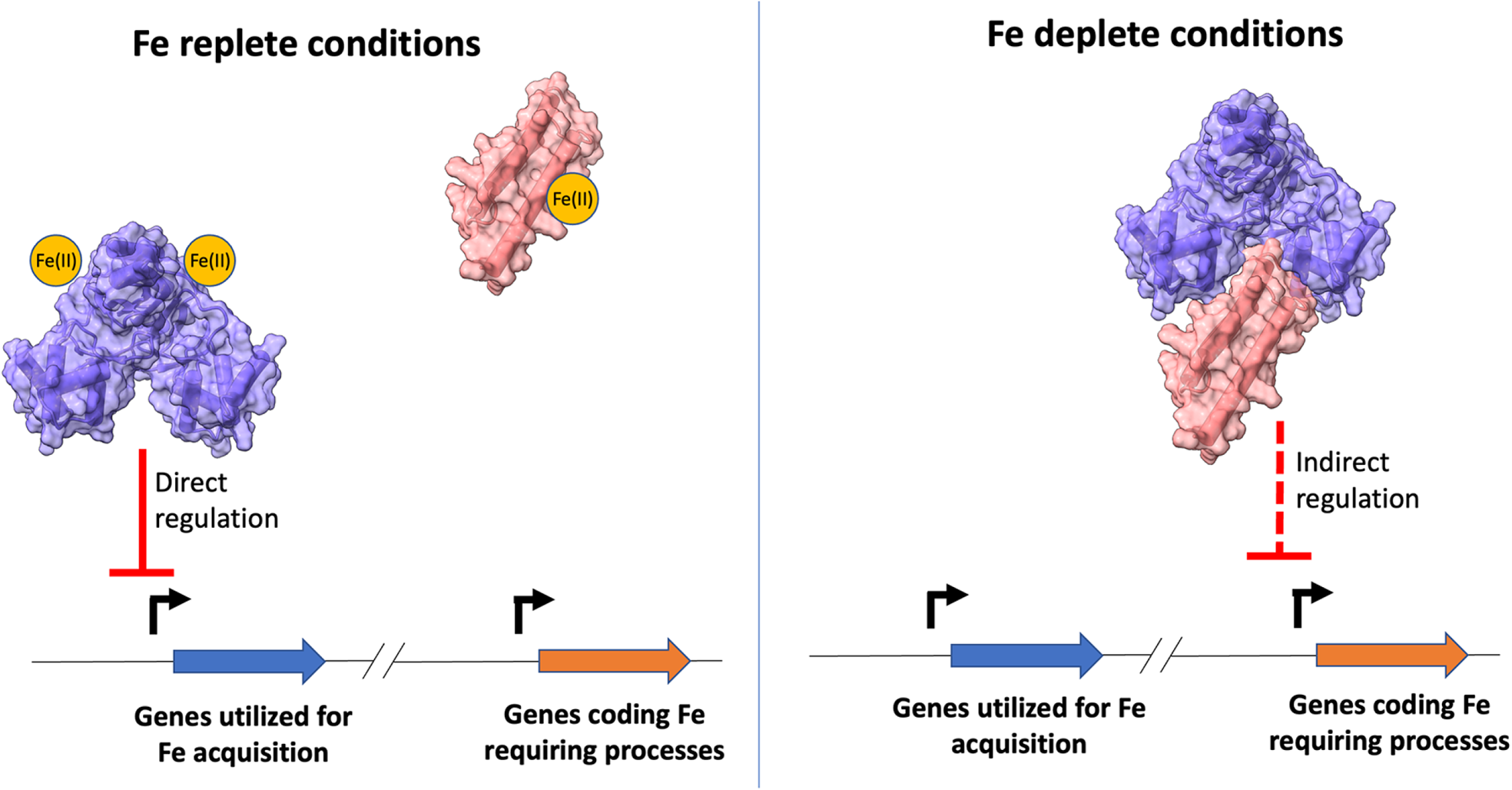
Working model for YlaN function. In our model, Fur (purple) and YlaN (pink) bind Fe(II) transiently. They are both metalated in Fe replete growth conditions and Fur directly prevents transcription of genes coding for Fe acquisition machinery. Upon Fe limitation, YlaN forms a complex with Fur and aids Fe(II) removal. Demetallated Fur has a decreased affinity for DNA and no longer acts as a transcriptional repressor. Fur derepression causes increased expression of genes coding for Fe acquisition systems and indirectly (broken line) decreases expression of genes coding for Fe requiring enzymes and processes.

YlaN also significantly interacted with Fe or FeS cluster-binding proteins. The genomes of *S. aureus* and *B. subtilis* lack the described bacterial Fe(II) binding proteins CyaY and IscX (40, 41), which act as Fe(II) chaperons and have been shown to interact with the Isc FeS cluster biosynthesis machinery (42). Further biochemical tests are required to determine how YlaN promotes Fur derepression and to determine whether YlaN can function as an Fe donor for FeS cluster synthesis or for the maturation of Fe proteins.

The findings herein have led to a model where YlaN acts as an Fe(II) chaperone. When the concentration of Fe(II) is sufficient, Fur and YlaN are metalated. Holo-Fur binds to operators to modulate transcription. When Fe is scarce, YlaN competes with Fur for Fe(II), and possibly DNA, resulting in demetallation of Fur and decreased DNA association. This results in derepression of Fe uptake systems and a reprogramming of central metabolism to compete under a low Fe environment.

## Materials and Methods

### Chemicals and growth conditions

All bacterial strains used in this study (Table 3) were derived from the community associated methicillin-resistant *Staphylococcus aureus* isolate USA300_LAC (43). Bacteria were grown at 37 °C in tryptic soy broth (TSB) (MP Biomedicals) and shaking at 200 rotations per minute. Solid media was generated by adding 1.5% (wt/vol) agar (VWR). Chelex-treated TSB was prepared as previously described (44). Unless stated otherwise, cells were cultured in 10 mL capacity culture tubes containing 1.5 mL of liquid medium. Quantitative growth was conducted using a 96-well plate reader BioTek 808E visible absorption spectrophotometer at 37 °C and shaking. For quantitative growth, bacteria strains were grown for 18 hours and washed twice with PBS before diluting them to an optical density at 600 nm (OD_600_) of 0.02 in 200 μL. Anaerobic growth was conducted using a 37 °C incubator inside of a COY anaerobic chamber containing an oxygen scavenging catalyst to maintain oxygen levels <1ppm. Anaerobic growth was quantified by measuring culture OD_600_ after 18 hours.

**Table 3.** Bacterial strains and plasmids used in this study.

| ***Staphylococcus aureus* USA300_LAC strains** |  |  |
| --- | --- | --- |
| **Strain name** | **genotype** | **Allele reference** |
| JMB1100 | USA300_LAC wild type (WT) | Alexander Horswill |
| JMB8689 | $\Delta ylaN::tetM$ | This study |
| JMB8428 | $ylaN::Tn$ | (89) this study |
| JMB1432 | $\Delta fur::tetM$ | (90) |
| JMB4536 | $hemB::Tn$ | (91) |
| JMB10678 | $\Delta ylaN::tetM fur^*$ | This study |
| JMB10675 | $proC::Tn$ | (89) and this study |
| JMB10677 | $\Delta ylaN::tetM proC::Tn$ | (89) and this study |
| JMB10676 | $fur^* proC::Tn$ | (89) and this study |
| JMB10678 | $\Delta ylaN::tetM fur^* proC::Tn$ | (89) and this study |
| JMB7525 | $fhuC::Tn$ | (6) |
| JMB10721 | $\Delta ylaN::tetM fhuC::Tn$ | (89) and this study |
| JMB10722 | $\Delta ylaN::tetM fur^* fhuC::Tn$ | (89) this study |
| JMB10641 | $ylaN::Tn \Delta fur::tetM$ | This study |
| ***Bacillus subtilis* strains** |  |  |
| BD630 | *hisA1 leuA8 metB5* | (92) |
| BD8592 | *hisA1 leuA8 metB5 P<sub>hyperspank</sub> yfp-ylaN* | This study |
| **Plasmids used in this study** |  |  |
| **Name** | **Function** | **Reference** |
| pEPSA5 | Inducible expression complementation | (93) |
| pEPSA5_ylaN | Inducible expression complementation | This study |
| pCM28 | Native promoter complementation | (94) |
| pCM28_ylaN | Native promoter complementation | This study |
| pJB38 | Mutant generation | (95) |
| pJB38_ΔylaN::tetM | Mutant generation | This study |
| pDR111-YFP | Protein interaction studies | (96) |
| pED2143 | Protein interaction studies |  |
| pOS_saeP_gfp | Transcriptional reporter | Shaun R. Brinsmade |
| pOS_isdC_gfp | Transcriptional reporter | This study |
| pOS_fhuA_gfp | Transcriptional reporter | This study |
| pOS_sbnA_gfp | Transcriptional reporter | This study |
| pOS-1-plgt | Genetic complementation *lgt* promoter | (97) |
| pOS-1-plgt_fur | Genetic complementation *lgt* promoter | (2) |

Streptonigrin sensitivity assays were performed as previously described (10). Overnight TSB cultures were washed with PBS and diluted to OD_600_ 0.1. One hundred μL of cell suspension were added to 4 mL of 3.5% TSA. For genetic complementation studies, the TSA top agar also contained 1% xylose and 30 μg mL^-1^ chloramphenicol (Cm). 5 μL of 2.5 mg mL^-1^ of streptonigrin diluted in DMSO was spotted atop the top agar overlay. The zone of inhibition was measured after overnight incubation at 37 °C.

When necessary, antibiotics were added at the final following concentrations: 100 μg mL^-1^ ampicillin (Amp); 10 μg mL^-1^ chloramphenicol (Cm); 10 μg mL^-1^ erythromycin (Erm); 3 μg mL^-1^ tetracycline (Tet). Protein concentrations were determined using Bradford reagent (Bio-Rad Laboratories Inc., Hercules, CA). DNA primers were purchased from IDT and are listed in Table S10. Molecular reagents were purchased from New England Biolabs, unless otherwise stated. Unless stated otherwise, all chemicals were purchased from Sigma-Aldrich (St. Louis, MO). Sanger DNA sequencing was performed at Azenta (South Plainfield, NJ).

### Plasmid and strain construction

*Escherichia coli* DH5α was used for plasmid preparation. The restriction minus strain *S. aureus* RN4220 was used for transformation (45) and transductions were carried out using bacteriophage 80α (46).

We used the pJB38 plasmid to create the Δ*ylaN::tetM* mutant as previously described (47). Briefly, the chromosomal regions upstream and downstream of *ylaN* were amplified by PCR using the following primer pairs: YCCylaNupFor and ylaNuptetRrev; tetRylaNdwnfor and ylaNdwnpJB38. We amplified *tetM* from strain JMB1432 using primers ylaNuptetRfor and tetRylaNdwnrev. The amplicons were gel purified and combined with pJB38 that had been digested with SalI and NheI. The fragments were combined using yeast recombinational cloning (48, 49). The pEPSA5_*ylaN* vector was constructed by amplifying *ylaN* from genomic DNA using the following primer pair: YlnA forEcoRI and YlnA revSalI. The pCM28_*ylaN* vector was made by amplifying *ylaN* from genomic DNA using the following primer pair: YlnA for5BamHI and YlnA revSalI. To create the pOS_*isdC_gfp* transcriptional reporter plasmid we amplified the *isdC* promoter using the following primer pair: pOS_saeP1_gfp with pOS_iscC_hindIII 5. For the pOS_*fhuA_gfp* transcriptional reporter plasmid we amplified the *fhuA* promoter using the following primer pair: pOS_fhuA_hindIII 5 and pOS_fhuA_kpnI 3. To create the pOS_tsr25_gfp transcriptional reporter, we amplified tsr25 using the following primer pair: HindIII tsr25p up and kpnI tsr25p dwn2. For the pOS_*sbnA_gfp* transcriptional reporter plasmid we amplified the *sbnA* promoter using the following primer pair: Sbn pro3 kpnI and Sbn pro5 hindIII. To create the pGEX-6P-1_*ylaN* expression vector we amplified *ylaN* from *S. aureus* using the following primer pair: ylaN5GSTBamHI and ylaN3XhoI. The amplicon was digested with BamHI and XhoI and ligated into similarly digested pGEX-6P-1. All plasmids were sequence verified.

To construct the *B. subtilis yfp-ylaN* construct, *ylaN* was amplified from *B. subtilis* BD630 genomic DNA (*hisA1 leuA8 metB5*), using the primers 5-yfp-ylaN and 3-yfp-ylaN. The vector pDR111-YFP was linearized by digestion with SalI and SphI. The In-Fusion HD cloning kit (Clontech) was used for cloning according to manufacturer’s instructions. The resulting plasmid was transformed into Stellar competent cells (Clontech), isolated, and verified by DNA sequence analysis performed by Eton Bioscience (Union, NJ). The resulting plasmid (pED2143) was transformed into BD630 selecting for spectinomycin resistance, placing the *yfp-ylaN* fusion under the control of the isopropyl β-D-1-thiogalactopyranoside (IPTG)-inducible *P_hyperspank_* promoter at the *amyE* locus, creating the strain BD8592. The strain was confirmed by sequencing performed by Eton Biosciences.

### Isolating and mapping suppressor mutations

Ten independent overnight cultures of the Δ*ylaN::tetM* strain were cultured overnight in TSB. Cultures were individually diluted 1:100 into phosphate buffered saline (PBS) and 100 μL was plated onto TSA plates with 700 μM DIP. One colony from each of the ten plates was taken and further characterized. The Δ*ylaN::tetM* mutants with suppressor mutations had growth defects indicative of the strains having a null mutation in *fur*. We amplified the *fur* allele from the Δ*ylaN::tetM* strain and each suppressor strain using the 1448 veri 5 and 1448 veri 3 primer pair. After gel purification, we sequenced the amplicons.

### Aconitase enzyme assay

The aconitase assays were performed as previously described (50) with slight modifications. *S. aureus* strains were grown in 1 mL of TSB with 0 or 120 μM DIP for four hours with shaking at 200 RPM. Cells were washed twice with PBS. A cell pellet was collected by centrifugation and then stored at −80 °C before thawing and assaying. Protein concentrations were determined using bicinchoninic acid (BCA) assay modified for a 96-well plate (51).

### RNA-seq analysis

*S. aureus* strains were cultured overnight in biological triplicates. Cultures were diluted into 2.5 mL of fresh TSB to an OD_600_ of 0.05 in 10 mL culture tubes with or without 120 μM DIP and subsequently cultured for eight hours aerobically with shaking. After eight hours, cultures were placed on ice, one mL of cells was harvested by centrifugation, washed with PBS, resuspended in 500 µL with RNA protect (QIAGEN), pelleted by centrifugation, and stored at −80°C. RNA extraction was performed as previously described (44). Cell pellets were thawed and washed twice with 0.5 mL of lysis buffer (20 mM RNase-free Sodium acetate, 1 mM EDTA, 0.5% SDS). The cells were lysed by the addition of 4 μg lysostaphin and incubated for 40 min at 37°C until confluent lysis was observed. RNA was isolated using TRIzol reagent according to the manufacturer’s instructions. RNA-seq libraries and DNA sequencing was conducted by SeqCenter (Pittsburg, PA).

RNA-seq data were submitted to GEO and the accession identifier GSE228225 was provided. RNA-seq data analysis was performed using CLC Genomics Workbench (Qiagen) as described previously (52). Raw data was imported and any remaining rRNA reads were filtered out using the “map reads to reference” function. RNAseq analysis was performed using the *S. aureus* USA300 FPR3757 genome updated to include annotations for sRNAs (53). Individual differential expression analyses were performed between biological triplicates and data normalized using quantile normalization. To eliminate lowly expressed genes, we excluded from further analysis genes in which the average expression values in both samples was less than 10 RPKM. To eliminate genes with ambiguously assigned reads we excluded from further analysis genes in which the percentage of assigned reads in any of the samples were less than 80% unique. Genes were considered significantly altered between two strains if they demonstrated a fold change >2 or <2 and a p-value <0.05 as determined by Students t-test. Transcripts exhibiting significant differences in expression were placed into functional categories based on their annotations in the COG database (54) as described previously (55). PCA plots were generated in CLC Genomics Workbench and exported to GraphPad Prism for visualization.

### RNA extraction, cDNA synthesis, and qPCR

*S. aureus* strains were cultured overnight in TSB and subsequently diluted in triplicate to an OD_600_ of 0.1 in 2.5 mL TSB with or without 500 μM DIP in 10 mL glass culture tubes. Cells were incubated at 37 °C with agitation for six hours after which time 1 mL cell pellets were treated with RNAprotect (Qiagen). Cell pellets were washed in 0.5 mL PBS pH 7.4, resuspended in 100 μL 50 mM tris pH 8 containing 6.7 μg lysostaphin, and then incubated at 37 °C with agitation for 30 minutes. The cell suspension was incubated at 65 °C for 5 minutes following the addition of 200 μL of 20 mM sodium acetate, 1 mM EDTA, 0.5% SDS with 13.4 μg lysostaphin. RNA isolation was performed as previously described (19). cDNA libraries were constructed using the High-Capacity cDNA Reverse Transcription kit (Biosystems). Quantitative real-time PCR was performed using an Applied Biosystems StepOnePlus thermal cycler. Data were analyzed using the comparative C_T_ method (44, 56).

### Metabolic modeling analyses

Model simulations were performed using the iYS854 genome-scale metabolic model of *S. aureus* (25) as previously described (57). RNA sequencing data were applied as modeling constraints using the iMAT algorithm (58, 59) using Gurobi and oxygen consumption rates were applied as additional modeling constraints for sampling in the COBRApy toolbox (60). The Fur deletion strain was modeled by setting the upper bounds for Fe(II) and Fe(III) to 0 to represent model uptake derepression. DIP treatment was modeling by creating sink reactions for Fe(II) and Fe(III) to represent Fe chelation. Simulations were performed by sampling each model 10,000 using optGpSampler (61).

### Metabolomic analyses

Strains were cultured overnight as biological triplicates before diluting into 2.5 mL of fresh TSB with or without 120 μM DIP to an OD_600_ of 0.05 in 10 mL culture tubes. Cells were subsequently cultured for eight hours aerobically with shaking or statically in an anaerobic chamber. After eight hours, OD_600_ was measured, and cultures were placed on ice. Samples for the metabolite profiling were prepared as described previously (62). Briefly, 1.5 mL of samples were collected, individual strains were normalized to the lowest OD_600_ of the triplicates, cells were pelleted, and washed twice with PBS. Cell pellets were resuspended in 1 mL of Methanol:Acetonitrile:Water (2:2:1) solution. Cells were lysed by bead beating (2 cycles, 40 s each, 6.0 m s^-1^) using a FastPrep homogenizer (MP Biomedicals) and 0.1-mm silica glass beads (MP Biomedicals). Samples were centrifuged twice at 14,500 rpm at 4 °C for 2 minutes and supernatant was retained. The supernatant was filtered with nylon membrane syringe filters (13 mm, 0.22 μm, Fisherbrand) and then samples were stored at −80 °C until metabolite analysis was performed.

Samples were analyzed at the metabolomics core of the Cancer Institute of New Jersey. HILIC separation was performed on a Vanquish Horizon UHPLC system (Thermo Fisher Scientific, Waltham, MA) with an XBridge BEH Amide column (150 mm × 2.1 mm, 2.5 μm particle size, Waters, Milford, MA) using a gradient of solvent A (95%:5% H_2_O:acetonitrile with 20 mM acetic acid, 40 mM ammonium hydroxide, pH 9.4) and solvent B (20%:80% H_2_O:acetonitrile with 20 mM acetic acid, 40 mM ammonium hydroxide, pH 9.4). The gradient was 0 min, 100% B; 3 min, 100% B; 3.2 min, 90% B; 6.2 min, 90% B; 6.5 min, 80% B; 10.5 min, 80% B; 10.7 min, 70% B; 13.5 min, 70% B; 13.7 min, 45% B; 16 min, 45% B; 16.5 min, 100% B; and 22 min, 100% B. The flow rate was 300 μL min^-1^. The column temperature was set to 25 °C. The autosampler temperature was set to 4 °C, and the injection volume was 5 μL. MS scans were obtained in negative mode with a resolution of 70,000 at m/z 200, in addition to an automatic gain control target of 3 x 106 and m/z scan range of 72 to 1000. Metabolite data was obtained using the MAVEN software package (63) (mass accuracy window: 5 ppm).

### CAS siderophore assay

Overnight cultures in TSB were diluted 100-fold into 1 mL of Chelex (Bio-Rad)- treated TSB with the addition of 25 μM zinc acetate, 25 μM MnCl_2_, 1 mM MgCl_2_ and 100 μM CaCl_2_ in 10 mL glass culture tubes. The cultures were incubated at 37 °C with shaking for 18 hours. The chrome azurol S siderophore assay was performed on the spent media using the modified microplate method as previously reported (64, 65).

### Whole cell metal quantification

*S. aureus* strains were grown for 18 hours overnight in TSB before diluting them to an OD of 0.05 (A_600_) in 7.5 mL of Chelex (Bio-Rad)-treated TSB in a 30 mL capacity culture tubes as described previously (44). Cells were allowed to grow with shaking for eight hours. Pre-weighted metal-free 15 mL propylene tubes were used to pellet the cells using a prechilled tabletop centrifuge (Eppendorf, Hauppauge, NY). Pellets were washed three times with 10 mL of ice-cold PBS. All samples were kept at −80 °C or on dry ice until processing.

Cell pellets were acid digested with 2 mL of Optima grade nitric acid (ThermoFisher, Waltham, MA) and 500 μL hydrogen peroxide (Sigma, St. Louis, MO) for 24 hr at 60°C. After digestion, 10 mL of UltraPure water (Invitrogen, Carlsbad, CA) was added to each sample. Elemental quantification on acid-digested liquid samples was performed using an Agilent 7700 inductively coupled plasma mass spectrometer (Agilent, Santa Clara, CA). The following settings were fixed for the analysis Cell Entrance = −40 V, Cell Exit = −60 V, Plate Bias = −60 V, OctP Bias = −18 V, and collision cell Helium Flow = 4.5 mL min^−1^. Optimal voltages for Extract 2, Omega Bias, Omega Lens, OctP RF, and Deflect were determined empirically before each sample set was analyzed. Element calibration curves were generated using ARISTAR ICP Standard Mix (VWR). Samples were introduced by a peristaltic pump with 0.5 mm internal diameter tubing through a MicroMist borosilicate glass nebulizer (Agilent). Samples were initially up taken at 0.5 rps for 30 s followed by 30 s at 0.1 rps to stabilize the signal. Samples were analyzed in Spectrum mode at 0.1 rps collecting three points across each peak and performing three replicates of 100 sweeps for each element analyzed. Sampling probe and tubing were rinsed for 20 s at 0.5 rps with 2% nitric acid between each sample. Data were acquired and analyzed using the Agilent Mass Hunter Workstation Software version A.01.02.

### Dioxygen consumption assay

*S. aureus* strains were grown overnight in 5 mL TSB in 30 mL culture tubes at 37 °C in a shaker (200 RPM). Strains were diluted 1:100 into 10 mL TSB containing 0, 250, 500 or 750 μM DIP in 125 mL flasks and incubated at 37 °C with agitation for four hours. The cultures were diluted to an OD_600_ of 0.025 in TSB containing 0, 250, 500 or 750 μM DIP prior to transfer of 200 μL to the wells of a Seahorse XF96 V3 PS cell culture microplate (Agilent). The seahorse XF sensor cartridge (Agilent) was hydrated in a non-CO_2_ incubator with sterile water overnight and equilibrated in XF calibrant (Agilent) for two hours prior to measurement. Measurements were taken for 15 cycles with a 3-minute mix and 3-minute measure cycle.

### Proton motive force measurement

Overnight cultures were grown in 2 mL TSB in 10 mL culture tubes at 37 °C with shaking. Strains were diluted 1:100 into 2 mL TSB +/− 500 μM DIP and incubated at 37 °C with shaking for six hours. Cell pellets were collected from 2 mL of culture and washed with 0.5 mL PBS, pH 7.4, and then resuspended in 0.5 mL PBS. The OD_600_ was adjusted to 0.085 in PBS. Thirty micromolar 3,3′-diethyloxacarbocyanine iodide (DiOC_2_(3)) +/− 5 μM carbonyl cyanide m-chlorophenyl hydrazone (CCCP) were added to the cell suspension followed by incubation at room temperature for 30 minutes. Fluorescence (excitation of 450 nm, emission of 670 nm) was measured using a Varioskan Lux plate reader (Thermo Scientific).

### Transcriptional reporter assays

Overnight cultures of *S. aureus* strains were grown overnight in 2 mL TSB supplemented with 10 μg mL^-1^ chloramphenicol in 10 mL culture tubes at 37 °C with shaking. *S. aureus* cultures were then subcultured 1:100 into 5 mL TSB supplemented with 10 μg mL^-1^ chloramphenicol +/− 500 μM DIP in 10 mL culture tubes and incubated at 37 °C with shaking for 8 hours. OD_600_ and GFP fluorescence (excitation 485 nm, emission 520 nm) were measured using a Varioskan Lux plate reader (Thermo Scientific). GFP fluorescence relative to OD_600_ was determined. The assay was performed in triplicate.

### Bioinformatic analyses

The predicted protein sequences from all assembled bacterial genomes designated as “reference” or “representative” in the NCBI database were retrieved on 03/14/2023. The *Staphylococcus aureus* YlaN sequence (ABD22267.1) was used as a query for a DIAMOND (v2.1.2; ‘--ultra-sensitive --max-target-seqs 0’) (66) search against the bacterial proteins retrieved from NCBI. Only hits with an *e*-value < 1e^-5^ were retained for downstream analysis; bacterial genomes that encode proteins with hits (*e*-value < 1e^-5^) to the YlaN query were considered YlaN-positive. Four Fur sequences (ABD21033.1, P54574, P0C6C8, P0A9A9), two from the species of interest in this study and two from species known to not encode YlaN, were used as queries for a DIAMOND (v2.1.2; ‘--ultra-sensitive --max-target-seqs 0’) search against the bacterial proteins retrieved from NCBI. Only hits with an *e*-value < 1e^-5^ were retained for downstream analysis. The hits from each of the four Fur query sequences were parsed separately, retaining the best (highest scoring) hit per genera. The top-scoring sequences (one per genera in the output) from each of the queries were combined and had redundant sequences (those identified by multiple queries) removed. The combined non-redundant sequences were aligned using mafft (v7.453; ‘--localpair --maxiterate 1000’) (67), with the resulting alignment used by fastme (v2.1.5) (68) for phylogenetic inference. Gotree (v0.4.4) and Goalign (v0.3.6) (69) were used for file format conversion between analysis steps and iTOL (70) was used for visualization. The filtering approach (down sampling to genera) applied to the Fur hits was performed to reduce the number of sequences to an amount that is computational tractable for phylogenetic analysis (i.e., it reduced the number of sequences in the resulting phylogeny from 10’s of thousands to only a few thousand).

The sequences that form the large clade of Firmicutes Fur sequences (Figure 7) were extracted for reanalysis. The protein sequences were realigned using mafft (’-- localpair --maxiterate 1000’) and had a phylogeny inferred using iqtree (v1.6.12; ‘-m MFP -bb 2000’). The alignment and phylogeny were compared together visually using iTOL. To improve readability a phylogeny with a reduced number of taxa (10% chosen at random from the full phylogeny) was also constructed and visualized using the same approach (Figure S11). A phylogeny of the Fur sequences from genomes which encode YlaN sequences (YlaN-positive genomes), and a phylogeny of the YlaN sequences from these genomes, were inferred from the Firmicutes extracted from Figure 7, with mafft and iqtree used for alignment and phylogeny inference (using the same versions and parameters as previously stated). Fur and YlaN sequences from genomes with multiple Fur or YlaN genes identified (using the previously described DIAMOND search results) were removed to prevent problems arising from paralogous sequences. The Fur and YlaN trees were compared using the tanglegram function (’sort=TRUE, rank_branches=TRUE’) from the dendextend (v1.17.1) R package (71).

To map changes in gene expression to the KEGG pathway maps, the predicted proteins for *Staphylococcus aureus* subsp. USA300_FPR3757 were downloaded from NCBI (NC_007793) and assigned KEGG Orthology (KO) numbers using the KEGG Automatic Annotation Server (gene dataset: *Staphylococcus aureus* USA300_FPR3757 [saa], search tool: BLAST) (72). The NCBI gene IDs were manually assigned to the old *Staphylococcus aureus* gene names used for expression analysis and used to transfer KO number annotations.

The structural model of *Staphylococcus aureus* Fur-YlaN complexes were built with ColabFold v.1.5.2: AlphaFold2 (Deep Mind) using MMseq2 (Max-Planck Institute for Biophysical Chemistry) (73) using the following amino acid sequences: ABD21033.1 and ABD22267.1. Five models were generated with three recycles. The final models were energy-minimized with Amber.

### YlaN interaction experiments

#### Immunopurification of YFP-YlaN

*B. subtilis* strain BD8592 was grown overnight in 50 mL of LB with 100 μg mL^-1^ spectinomycin. After incubation, cells were diluted 1:100 (v/v) into 2 L of fresh LB media with the addition of 0.5 mM isopropyl-β-d-1-thiogalactosidase (IPTG) and grown for 1.5 hours for exponential phase or 4 hours for stationary phase. Growth was monitored hourly by the optical density at OD_600_. Cells were harvested by centrifugation, frozen as pellets in liquid nitrogen, and subjected to cryogenic cell lysis as described previously (27). 0.75 g of frozen cell powder was immediately added to 10 mL of lysis buffer (20 mM HEPES, pH 7.4, 100 mM potassium acetate, 2 mM MgCl_2_, 0.1% tween-20 (v/v), 1 µM ZnCl_2_, 1 µM CaCl_2_, 0.25% Triton-X, 200 mM NaCl, 1:100 protease inhibitor cocktail (Sigma) and 0.1 mg mL^-1^ phenylmethylsulphonyl fluoride (PMSF)). The resulting suspension was homogenized for 20 s using a PT 10-35 GT Polytron (Kinematica) and centrifuged for 10 min at 8000 x *g* at 4°C. The soluble fraction was mixed with 10 µL of GFP-Trap magnetic agarose (Proteintech) for 1 h with gentile rotation at 4°C. The magnetic beads were recovered and washed four times with lysis buffer without inhibitors and two times with PBS. Proteins were eluted directly into 50 µL of TEL buffer (106 mM Tris-HCl, 141 mM Tris-base, 0.5 mM EDTA, 2.0% LDS, pH 8.5) for in-solution digestion (74, 75).

#### Preparation of samples for mass spectrometry

Samples were alkylated and reduced with final concentrations of 30 mM chloroacetamide and 10 mM Tris(2-carboxyethyl)phosphine hydrochloride (TCEP, pH 7.0), and heated to 95 °C for 5 minutes. Protein digestion was performed using a filter-aided sample preparation method (FASP), (76–78). Briefly, filters [Vivacon 500 centrifugal filters (10K cut-off), Sartorius Stedim Biotech, Goettingen, Germany] were used for desalting and overnight trypsin digestion at 37°C with a 1:50 enzyme to protein ratio in 40 mM HEPES, pH 7.4, 0.1% sodium deoxycholate. Following digestion, the detergent was removed from the samples through acidification with 1% triflouroacetic acid (TFA, final concentration) as described (79, 80). Soluble peptides were desalted using SDB-RPS Stage Tips and eluted in 50 mL of 5% ammonium hydroxide-80% acetonitrile (ACN) (81). Samples were evaporated to near dryness by vacuum centrifugation and resuspended in 1% formic acid (FA)/4% ACN to bring the total volume to 10 mL.

#### Mass spectrometry

Samples were analyzed by nano-liquid chromatography-tandem mass spectrometry (nLC-MS/MS) on a Dionex Ultimate 3000 RSLC coupled directly to an LTQ-Orbitrap Velos mass spectrometer (ThermoFisher Scientific, San Jose, CA), through a Nanospray Flex ion source (ThermoFisher Scientific). Approximately 4 µL of peptides were directly injected for analysis. Instrument parameters and settings were as described previously (82), except peptides were separated using a 150 min linear reverse phase gradient. The mass spectrometer was operated in a data-dependent acquisition mode with each cycle of analysis containing a single full-scan mass spectrum (*m*/*z* = 350–1700) in the orbitrap (*r* = 120,000 at *m*/*z* = 400) followed by collision-induced dissociation (CID) MS/MS of the top 15 most abundant ions, with dynamic exclusion enabled.

#### Informatics workflow

The MS/MS spectra were extracted using the Proteome Discoverer software platform (ver. 1.4, Thermofisher Scientific). All spectra were subsequently analyzed using SEQUEST (ver. 1.4.0.288. ThermoFisher Scientific) and X! Tandem (ver. CYCLONE (2010.12.01.1), The GPM, thegpm.org) for database searching against the UniProt SwissProt sequence database (downloaded 04/2017) comprised of *Bacillus subtilis* and *Escherichia coli* reference proteome sequences, including common contaminant sequences (total of 8604 sequences). X! Tandem was set up to search a reverse concatenated subset of the same database. The processing workflow parameters were defined as follows: peptides with at least 6 amino acids, full trypsin cleavage specificity, and up to 2 missed cleavages. Database searching was performed with precursor and fragment ion mass tolerances of 10 parts per million (ppm) and 0.4 Da, respectively. Cysteine carbamidomethylation was denoted as a fixed modification and methionine oxidation, asparagine deamidation, phosphorylation of serine and threonine, lysine acetylation, and n-terminal acetylation, as variable modifications. Peptide and protein identifications were validated using Scaffold (ver. 4.8.1, Proteome Software, Inc.), with a peptide and protein false discovery rate (FDR) threshold of < 1.0%. For label-free quantification, the intensity based on the sum of the three highest intensity peptides for each protein (T3PQ) was used (83).

### YlaN purification

*E. coli* strain BL21(DE3) containing pGEX-6P-1_*ylaN* was cultured in 3 L of 2X LB supplemented with ampicillin (50 μg mL^-1^) at 30°C with shaking. After reaching an OD_600_ of 0.5, the temperature was shifted to ∼25°C and *ylaN* expression was induced with 100 μM IPTG for ∼16 hrs. The cells were harvested by centrifugation and resuspended in purification buffer (150 mM NaCl, 50 mM Tris-HCl, pH 7.5) containing 1% weight per volume lysozyme. Cells were disrupted using a French pressure cell and cell debris removed by centrifugation. The cell free extract was filtered through a 0.45 μM filter before loading into a 5 mL glutathione Sepharose 4 fast flow column (Cytiva).

The GST column was equilibrated with purification buffer and filtered extract was loaded onto the column. The column was washed with 25 column volumes of purification buffer. After washing, the column was incubated with 1 column volume of cleavage buffer (50 mM Tris-HCl, pH 7.0, 150 mM NaCl, 1 mM EDTA, 1 mM dithiothreitol containing PreScission Protease (Cytiva). Following overnight incubation, YlaN was eluted using 5 mL of cleavage buffer. Eluted protein was concentrated using spin concentrators (Millipore Amicon Ultra-15 -3K) to ∼1.5-2 mL and subsequently dialyzed in one liter of storage buffer (150 mM NaCl, 50 mM Tris-HCl, pH 7.5, 5% (v/v) glycerol) three individual times to remove EDTA. YlaN was flash frozen and stored at −80°C until use. Protein concentration was measured using a BCA assay.

### Fe-Binding Competition Assay

The ferrous ion binding affinity of YlaN was measured as previously reported (29). Briefly, this assay exploits Mag-Fura-2 (Molecular Probes) as a metal chelator chromophore to measure the binding of ferrous ions to an additional biomolecule (i.e., YlaN) under complex heterogeneous conditions. Mag-Fura-2 forms a 1:1 complex with Fe(II) and displays a maximum absorbance at 325 nm when metal is bound; this is in contrast to a feature at 365 nm when the chelator is in the apo state (29, 30, 32). The transition of the absorbance feature at 365 nm, observed during metal loading of YlaN, was measured using a Shimadzu UV-1800 spectrophotometer housed within a Coy anaerobic chamber. Titration data for adding a divalent metal ion into the protein/Mag-Fura-2 mixture were collected anaerobically at room temperature using a 1 cm quartz cuvette. All spectra were collected anaerobically in 20 mM Tris, 150 mM NaCl (pH 7.5) buffer.

Samples were prepared for analysis in the following manner. Reagents/buffers were Ar(g) purged and equilibrated overnight within a Coy anaerobic wet chamber prior to experimentation. Holo- and apo-protein solution samples were prepared immediately before use under the same buffer and maintained under anaerobic conditions. Independent protein samples, incubated in 5 mM TCEP prior to titrations, were dialyzed using anaerobic buffer before analysis to remove the TCEP. Mag-Fura-2 concentrations were varied between 1 and 4 µM, while the protein concentration was held constant at 2 µM. A solution of 2 mM ammonium ferrous sulfate hexahydrate, prepared in the anaerobic buffer listed previously was added in progressive increments until absorption saturation was reached. After each addition of aqueous ferrous ions, an absorption spectrum was collected between the wavelength of 200-800 nm. Initial apo Mag-Fura-2 concentrations were determined using the molar absorptivity (ε) value of 29,900 M^-1^cm^-1^ for the compound, measured at the wavelength of 365 nm (28). The absorbance at 365 nm, corrected for dilution, was then used to calculate binding parameters. Binding data were simulated with the program DYNAFIT (84), using a non-linear least squares analysis script to identify the binding capacity and metal stoichiometry in a manner previously outlined (28, 30). Each titration experiment was simulated using both one and two-site metal binding models.

### Circular Dichroism Spectroscopy

Apo- and holo-YlaN samples were characterized for secondary structure content using a Jasco 1500 spectrophotometer as described previously (30). Briefly, samples were prepared anaerobically in a Coy wet chamber, loaded into a 0.1 cm quartz cuvette, then transferred to the nitrogen purged sample chamber of the instrument where data was collected at 27 °C. Spectra were collected on 5 - 10 µM protein concentration samples 5 mM NaPO_4_ buffer solutions at pH 7.5. For reproducibility, an average of thirty scans were collected on three samples per state and results were averaged for final analysis. Before spectral collection, a baseline at each wavelength was subtracted to eliminate buffer signal. Data were analyzed using the Jasco CD Pro analysis software and simulated using the CONTIN method including the SP29, SP37, SP43, SMP50 and SMP56 reference sets (31). Values obtained from simulations using each database were averaged to obtain final analysis parameters.

### Differential Scanning Calorimetry

Thermal stability parameters for apo- and holo-YlaN were evaluated using a VP Differential Scanning Calorimeter (TA Instruments) housed anaerobically in a Coy wet chamber. Protein and iron solutions were prepared anaerobically in 20 mM Tris, 150 mM NaCl solution at a pH of 7.5. Buffer was independently degassed with argon prior to sample preparation and loading. Protein concentrations between 3-5 mg mL^-1^ were used, and a 2.0 mM ferrous ion solution was added to obtain samples up to a 1:1 protein to Fe ratio at a final volume of 600 μL. DSC scans were run at a rate of 1 ℃ per minute over a temperature range of 10-90 °C. Pressure was maintained above 2.95 atm throughout the scan. Data were analyzed using the Nano Analyze software provided by TA Instruments. Baseline subtraction of buffer alone or buffer with iron at concentrations that matched the protein samples were measured as controls. Peak deconvolution of the protein spectra was achieved using the two-state scaled mathematical model that incorporates the A_w_ factor, which accounts for inaccuracies in protein concentration upon denaturation. Data presented represents an average of three sample results.

### Ray Absorption Spectroscopy

Fe(II)-YlaN samples were prepared in 20 mM Tris and 150 mM NaCl. The sample was brought to a 30% glycerol concentration for cryo-protection, and samples were flash frozen and stored in liquid nitrogen immediately after loading into prewrapped 2 mm Leucite XAS cells until exposed to the beam. Fe XAS was collected at the Stanford Synchrotron Radiation Lightsource (SSRL) on beamline 7-3. This beamline is equipped with a Si[220] double crystal monochromator with a mirror present upstream for focusing and harmonic rejection. We used a Canberra 30 element germanium solid state detector to measure protein fluorescence excitation spectra. Temperature during collection was maintained at 10 K by an Oxford Instruments continuous-flow liquid helium cryostat and data was collected with the solar slits utilizing a 3 µm Mn filter placed between the cryostat and detector to diminish background scattering. XAS spectra were collected in 5 eV increments in the pre-edge region, 0.25 eV increments in the edge region, and 0.05 Å^-1^ increments in the EXAFS region to k = 14 Å^-1^, integrated from 1 to 25 s in a k^3^-weighted manner for a total scan length of ∼40 min. Fe foil absorption spectra were simultaneously collected with each respective protein run, and used for calibration and assigning first inflection points for the metals (7,111.2eV)

XAS spectra were processed and analyzed using the EXAFSPAK program suite written for Macintosh OS-X (85), integrated with the Feff v8 software (86) for theoretical model generation. Normalized XANES data was subjected to edge analysis for both metals and in the case of Fe, to pre-edge analysis as well. The Fe 1s-3d pre-edge peak analysis was completed as described previously (87); peak area was determined over the energy range of 7,110–7,116 eV. Oxidation state was deduced from the first inflection energies of the respective edges (32). Data were collected to k = 14 Å^-1^, which corresponds to a spectral resolution of 0.121 Å^-1^ for all metal–ligand interactions (32); therefore, only independent scattering environments at distances >0.121 Å were considered resolvable in the EXAFS fitting analysis. Data were fit using both single and multiple scattering model amplitudes and phase functions to simulate Fe-O/N, -S, and - Fe ligand interactions. During Fe data simulations, a scale factor (Sc) of 0.95 and threshold shift (ΔE_0_) value of −10 eV (Fe–O/N/C), −12 eV (Fe–S) and −15 eV (Fe–Fe) were used. These values were obtained from fitting crystallographically characterized small molecule Fe (87). The best-fit EXAFS simulations were based on the lowest mean square deviation between data and fit, corrected for the number of degrees of freedom (*F’*) (88). During the standard criteria simulations, only the bond length and Debye-Waller factor were allowed to vary for each ligand environment.

## Supplemental Figure legends

**Figure S1.** Growth of the wild type and Δ*ylaN::tetM* strains in TSB medium. Overnight cultures of the wild type (JMB1100) and the *ylaN::tetM* mutant (JMB8689) were diluted 1:1000 into 5 mL of fresh TSB in 25 mL capacity culture tube. Cells were cultured at 37 °C with shaking at 200 rpm.

**Figure S2.** The ^57^Fe load was quantified in whole cells using ICPMS after culture in TSB medium. The ratio of ^57^Fe / ^31^P is displayed for the *proC::Tn* (JMB10675), Δ*ylaN::tetM proC::Tn* (JMB10677), and *fur* proC::Tn* (JMB10676), and Δ*ylaN::tetM fur* proC::Tn* (JMB10678) strains. Student’s t-tests were performed on the data and * indicates p < 0.05.

**Figure S3.** Growth of the WT (JMB1100), Δ*ylaN::tetM* (JMB8689), Δ*ylaN::tetM fur\** (JMB10678), *fhuC::Tn* (JMB7525), Δ*ylaN::tetM fhuC::Tn* (JMB10721), Δ*ylaN::tetM fur* fhuC::Tn* (JMB10722) strains with or without 2,2-dipyridyl (DIP) or EDDHA. Overnight cultures were grown in TSB, serial diluted, and spot plated. Pictures of a representative experiment are displayed.

**Figure S4.** PCA plot of significantly altered RNA abundances in the wild type (WT; JMB1100) and Δ*fur::tetM* (JMB1432) strains after culture in TSB media with and without 150 μM 2,2-dipyridyl. RNAs were isolated from biological triplicates.

**Figure S5.** A plot of the Log2 fold change determined by RNA sequencing was plotted against the Log2 fold change determined using qPCR.

**Figure S6.** A Venn diagram displaying the transcriptional overlap between the WT (JMB1100) vs Δ*fur::tetM* (JMB1432) regulon and the WT vs WT + 150 μM 2,2-dipyridyl (DIP) regulon.

**Figure S7.** The abundances of select RNAs of the WT (JMB1100) and Δ*fur::tetM* (JMB1432) after growth in TSB with and without 150 μM 2,2-dipyridyl (DIP). Normalized RNA reads were from RNA sequencing data.

**Figure S8.** Transcripts exhibiting significant differences in expression in the WT (JMB1100) vs Δ*fur::tetM* (JMB1432) (Panel A) and the WT vs WT + 150 μM 2,2-dipyridyl (DIP) (Panel B) regulons were placed into functional categories based on their annotations in the COG database.

**Figure S9.** A *hemB::Tn* mutant is defective in respiration and generating a proton motive force. **Panel A:** Dioxygen consumption rates (OCR) of the WT (JMB1100) and *hemB::Tn* (JMB4536) strains cultured in TSB medium. **Panel B:** The membrane potentials of the WT and *hemB::Tn* strains were measured using the fluorescent dye 3′3′-diethyloxacarbocyanine iodide (DiOC_2_). Overnight cultures were grown in tryptic soy broth (TSB). Cells were washed with phosphate buffered saline (PBS) pH 7.4 and then resuspended in 0.5 mL PBS pH 7.4. Thirty micromolar 3,3′-diethyloxacarbocyanine iodide (DiOC_2_(3)) +/− 5 μM carbonyl cyanide m-chlorophenyl hydrazone (CCCP) were added to the cell suspensions followed by incubation at room temperature for 30 minutes. Fluorescence (excitation of 450 nm, emission of 670 nm) was measure. All data represent the average of biological triplicates and standard deviations are displayed. Student’s t-tests were performed on the data and * indicates p < 0.05.

**Figure S10.** The abundances of RNAs from the WT (JMB1100) and Δ*fur::tetM* (JMB1432) strains were determined using quantitative PCR. Fold changes were determined using the ΔΔCt method. Student’s t-tests were performed on the data and * indicates p < 0.05.

**Figure S11:** Down sampled phylogeny and associated alignment of Firmicutes Fur sequences. Sequences were randomly chosen from those in the Firmicutes Fur clade (Figure 7) to facilitate visualization of the relationship (tree shown on left side of the figure) between the YlaN-positive (black bars in the middle of the figure) and -negative sequences and their associated aligned protein sequences (left side of the figure; residues colored using the clustal scheme). Phylogeny were constructed using maximum-likelihood with automatic model selection and node support tested *via* 2,000 ultrafast phylogenetic bootstraps.

**Figure S12:** Tanglegram of Firmicutes Fur and YlaN proteins. Phylogenies of Firmicutes Fur (left) and YlaN (right) proteins from YlaN-positive genomes. Lines are used to connect the same tips in each tree, with groups of tips from clades shared between both trees shown using colored lines. Phylogenies were constructed using maximum-likelihood with automatic model selection and node support tested *via* 2,000 ultrafast phylogenetic bootstraps. Branch lengths have been scaled and only nodes with ≥ 95% bootstrap support are annotated to improve readability.

**Figure S13.** AlphaFold2 structural models of the Fur-YlaN complex. **Panel A:** Surface representation of the complex modeled in one-to-one stoichiometry. YlaN (ABD22267.1) is displayed in pink and Fur (ABD21033.1) in shown in purple. Some of the conserved residues of YlaN helix two are highlighted in yellow. **Panel B:** Surface representation of the Fur-YlaN complex modeled in two to one stoichiometry. **Panel C:** The predicted Local Distance Difference Test (pLDDT) values calculated for five AlphaFold2 ranked models. The “rank 1” model of the Fur-YlaN complex formed in one-to-one stoichiometry is presented in Panel A. **Panel D:** The Local Distance Difference Test (lDDT) values calculated for five AlphaFold2 ranked models. The “rank 1” model of the Fur-YlaN complex formed in two to one stoichiometry is presented in Panel B. The accuracy of all AlphaFold2 predictions is reflected by high pLDDT values, in particular those obtained for Fur.

## Table Legends

**Table S2.** Comparisons in the abundances of RNAs isolated from the WT (JMB1100) and Δ*fur::tetM* (JMB1432) strains after culture in TSB medium with and without 120 μM 2,2-dipyridyl (DIP).

**Table S3.** Metabolic modeling simulations corresponding to WT, DIP, Fur, and FUR_DIP strains using the iYS854 genome-scale metabolic model for *S. aureus* (PMID: 30625152).

**Table S4**. Concentrations of metabolites isolated from the WT (JMB1100) and Δ*fur::tetM* (JMB1432) strains after culture in TSB medium with and without 120 μM 2,2-dipyridyl (DIP). The strains were incubated either aerobically or anaerobically in the absence of a terminal electron acceptor to force fermentative growth. Metabolite concentrations were standardized by divided them by the optical density of the culture at the time of harvest.

**Table S5.** List of identified proteins from the YFP-YlaN immunopurifications performed during logarithmic (LOG) and stationary (STAT) phase growth, with at least 4 identified peptides in one growth phase and an FDR < 1.0%. The number of unique identified peptides for each protein are displayed. The MS intensity represents the sum of the three highest intensity peptides for each protein. Proteins in red text represent those that bind iron, contain FeS clusters, or are part of the FeS cluster biosynthesis machinery.

## REFERENCES

1. Imlay JA, Chin SM, Linn S. 1988. Toxic DNA damage by hydrogen peroxide through the Fenton reaction in vivo and in vitro. Science 240:640–2.

2. Torres VJ, Attia AS, Mason WJ, Hood MI, Corbin BD, Beasley FC, Anderson KL, Stauff DL, McDonald WH, Zimmerman LJ, Friedman DB, Heinrichs DE, Dunman PM, Skaar EP. 2010. *Staphylococcus aureus* Fur regulates the expression of virulence factors that contribute to the pathogenesis of pneumonia. Infect Immun 78:1618–28.

3. Hammer ND, Skaar EP. 2011. Molecular mechanisms of *Staphylococcus aureus* iron acquisition. Annu Rev Microbiol 65:129–47.

4. Grim KP, San Francisco B, Radin JN, Brazel EB, Kelliher JL, Parraga Solorzano PK, Kim PC, McDevitt CA, Kehl-Fie TE. 2017. The Metallophore Staphylopine Enables *Staphylococcus aureus* To Compete with the Host for Zinc and Overcome Nutritional Immunity. mBio 8.

5. Ghssein G, Brutesco C, Ouerdane L, Fojcik C, Izaute A, Wang S, Hajjar C, Lobinski R, Lemaire D, Richaud P, Voulhoux R, Espaillat A, Cava F, Pignol D, Borezee-Durant E, Arnoux P. 2016. Biosynthesis of a broad-spectrum nicotianamine-like metallophore in *Staphylococcus aureus*. Science 352:1105–9.

6. Roberts CA, Al-Tameemi HM, Mashruwala AA, Rosario-Cruz Z, Chauhan U, Sause WE, Torres VJ, Belden WJ, Boyd JM. 2017. The Suf Iron-Sulfur Cluster Biosynthetic System Is Essential in *Staphylococcus aureus*, and Decreased Suf Function Results in Global Metabolic Defects and Reduced Survival in Human Neutrophils. Infect Immun 85.

7. Steingard CH, Helmann JD. 2023. Meddling with Metal Sensors: Fur-Family Proteins as Signaling Hubs. J Bacteriol 205:e0002223.

8. Price EE, Boyd JM. 2020. Genetic Regulation of Metal Ion Homeostasis in Staphylococcus aureus. Trends Microbiol 28:821–831.

9. Troxell B, Hassan HM. 2013. Transcriptional regulation by Ferric Uptake Regulator (Fur) in pathogenic bacteria. Front Cell Infect Microbiol 3:59.

10. Bagg A, Neilands JB. 1987. Ferric uptake regulation protein acts as a repressor, employing iron (II) as a cofactor to bind the operator of an iron transport operon in Escherichia coli. Biochemistry 26:5471–7.

11. Escolar L, de Lorenzo V, Perez-Martin J. 1997. Metalloregulation in vitro of the aerobactin promoter of Escherichia coli by the Fur (ferric uptake regulation) protein. Mol Microbiol 26:799–808.

12. Coronel-Tellez RH, Pospiech M, Barrault M, Liu W, Bordeau V, Vasnier C, Felden B, Sargueil B, Bouloc P. 2022. sRNA-controlled iron sparing response in Staphylococci. Nucleic Acids Res 50:8529–8546.

13. Peters JM, Colavin A, Shi H, Czarny TL, Larson MH, Wong S, Hawkins JS, Lu CH, Koo BM, Marta E, Shiver AL, Whitehead EH, Weissman JS, Brown ED, Qi LS, Huang KC, Gross CA. 2016. A Comprehensive, CRISPR-based Functional Analysis of Essential Genes in Bacteria. Cell 165:1493–506.

14. Rauen U, Springer A, Weisheit D, Petrat F, Korth HG, de Groot H, Sustmann R. 2007. Assessment of chelatable mitochondrial iron by using mitochondrion-selective fluorescent iron indicators with different iron-binding affinities. Chembiochem 8:341–52.

15. Yunta F, Garcia-Marco S, Lucena JJ, Gomez-Gallego M, Alcazar R, Sierra MA. 2003. Chelating agents related to ethylenediamine bis(2-hydroxyphenyl)acetic acid (EDDHA): synthesis, characterization, and equilibrium studies of the free ligands and their Mg2+, Ca2+, Cu2+, and Fe3+ chelates. Inorg Chem 42:5412–21.

16. Bergan T, Klaveness J, Aasen AJ. 2001. Chelating agents. Chemotherapy 47:10–4.

17. Luo Y, Han Z, Chin SM, Linn S. 1994. Three chemically distinct types of oxidants formed by iron-mediated Fenton reactions in the presence of DNA. Proc Natl Acad Sci U S A 91:12438–42.

18. DeGraff W, Hahn SM, Mitchell JB, Krishna MC. 1994. Free radical modes of cytotoxicity of adriamycin and streptonigrin. Biochem Pharmacol 48:1427–35.

19. Mashruwala AA, Pang YY, Rosario-Cruz Z, Chahal HK, Benson MA, Mike LA, Skaar EP, Torres VJ, Nauseef WM, Boyd JM. 2015. Nfu facilitates the maturation of iron-sulfur proteins and participates in virulence in *Staphylococcus aureus*. Mol Microbiol 95:383–409.

20. Sheldon JR, Heinrichs DE. 2012. The iron-regulated staphylococcal lipoproteins. Front Cell Infect Microbiol 2:41.

21. Speziali CD, Dale SE, Henderson JA, Vines ED, Heinrichs DE. 2006. Requirement of Staphylococcus aureus ATP-binding cassette-ATPase FhuC for iron-restricted growth and evidence that it functions with more than one iron transporter. J Bacteriol 188:2048–55.

22. Hammer ND, Reniere ML, Cassat JE, Zhang Y, Hirsch AO, Indriati Hood M, Skaar EP. 2013. Two heme-dependent terminal oxidases power *Staphylococcus aureus* organ-specific colonization of the vertebrate host. MBio 4.

23. Sims PJ, Waggoner AS, Wang CH, Hoffman JF. 1974. Studies on the mechanism by which cyanine dyes measure membrane potential in red blood cells and phosphatidylcholine vesicles. Biochemistry 13:3315–30.

24. Bryan LE, Kwan S. 1983. Roles of ribosomal binding, membrane potential, and electron transport in bacterial uptake of streptomycin and gentamicin. Antimicrob Agents Chemother 23:835–45.

25. Seif Y, Monk JM, Mih N, Tsunemoto H, Poudel S, Zuniga C, Broddrick J, Zengler K, Palsson BO. 2019. A computational knowledge-base elucidates the response of Staphylococcus aureus to different media types. PLoS Comput Biol 15:e1006644.

26. Cheung J, Beasley FC, Liu S, Lajoie GA, Heinrichs DE. 2009. Molecular characterization of staphyloferrin B biosynthesis in Staphylococcus aureus. Mol Microbiol 74:594–608.

27. Carabetta VJ, Tanner AW, Greco TM, Defrancesco M, Cristea IM, Dubnau D. 2013. A complex of YlbF, YmcA and YaaT regulates sporulation, competence and biofilm formation by accelerating the phosphorylation of Spo0A. Molecular Microbiology 88:283–300.

28. Rodrigues AV, Kandegedara A, Rotondo JA, Dancis A, Stemmler TL. 2015. Iron loading site on the Fe-S cluster assembly scaffold protein is distinct from the active site. Biometals 28:567–76.

29. Koebke KJ, Batelu S, Kandegedara A, Smith SR, Stemmler TL. 2020. Refinement of protein Fe(II) binding characteristics utilizing a competition assay exploiting small molecule ferrous chelators. J Inorg Biochem 203:110882.

30. Dzul SP, Rocha AG, Rawat S, Kandegedara A, Kusowski A, Pain J, Murari A, Pain D, Dancis A, Stemmler TL. 2017. In vitro characterization of a novel Isu homologue from Drosophila melanogaster for de novo FeS-cluster formation. Metallomics 9:48–60.

31. Sreerama N, Woody RW. 2004. Computation and analysis of protein circular dichroism spectra. Methods Enzymol 383:318–51.

32. Bencze KZ, Kondapalli KC, Stemmler TL. 2007. X-Ray Absorption Spectroscopy, p 513–28. In Scott RA, Lukehart CM (ed), Applications of Physical Methods in Inorganic and Bioinorganic Chemistry: Handbook, Encyclopedia of Inorganic Chemistry, 2nd Edition. John Wiley & Sons,LTD, Chichester, UK.

33. Nakano MM, Dailly YP, Zuber P, Clark DP. 1997. Characterization of anaerobic fermentative growth of Bacillus subtilis: identification of fermentation end products and genes required for growth. J Bacteriol 179:6749–55.

34. Cruz Ramos H, Hoffmann T, Marino M, Nedjari H, Presecan-Siedel E, Dreesen O, Glaser P, Jahn D. 2000. Fermentative metabolism of Bacillus subtilis: physiology and regulation of gene expression. J Bacteriol 182:3072–80.

35. Py B, Barras F. 2010. Building Fe-S proteins: bacterial strategies. Nat Rev Microbiol 8:436–46.

36. Ledala N, Zhang B, Seravalli J, Powers R, Somerville GA. 2014. Influence of iron and aeration on Staphylococcus aureus growth, metabolism, and transcription. J Bacteriol 196:2178–89.

37. Friedman DB, Stauff DL, Pishchany G, Whitwell CW, Torres VJ, Skaar EP. 2006. *Staphylococcus aureus* redirects central metabolism to increase iron availability. PLoS Pathog 2:e87.

38. Nunoshiba T, Obata F, Boss AC, Oikawa S, Mori T, Kawanishi S, Yamamoto K. 1999. Role of iron and superoxide for generation of hydroxyl radical, oxidative DNA lesions, and mutagenesis in Escherichia coli. J Biol Chem 274:34832–7.

39. L. D, R. B, K. S, B. H, R. W, J. R, J. S. 2023. Control of iron homeostasis by a regulatory protein-protein interaction in Bacillus subtilis: The FurA (YlaN) acts as an antirepressor to the ferric uptake regulator Fur. in review.

40. Bou-Abdallah F, Adinolfi S, Pastore A, Laue TM, Dennis Chasteen N. 2004. Iron binding and oxidation kinetics in frataxin CyaY of *Escherichia coli*. Journal of Molecular Biology 341:605–615.

41. Pastore C, Adinolfi S, Huynen MA, Rybin V, Martin S, Mayer M, Bukau B, Pastore A. 2006. YfhJ, a molecular adaptor in iron-sulfur cluster formation or a frataxin-like protein? Structure 14:857–67.

42. Roche B, Huguenot A, Barras F, Py B. 2015. The iron-binding CyaY and IscX proteins assist the ISC-catalyzed Fe-S biogenesis in *Escherichia coli*. Mol Microbiol 95:605–23.

43. Pang YY, Schwartz J, Bloomberg S, Boyd JM, Horswill AR, Nauseef WM. 2013. Methionine Sulfoxide Reductases Protect against Oxidative Stress in *Staphylococcus aureus* Encountering Exogenous Oxidants and Human Neutrophils. J Innate Immun doi:10.1159/000355915.

44. Al-Tameemi H, Beavers WN, Norambuena J, Skaar EP, Boyd JM. 2021. Staphylococcus aureus lacking a functional MntABC manganese import system has increased resistance to copper. Mol Microbiol 115:554–573.

45. Kreiswirth BN, Löfdahl S, Betley MJ, O’Reilly M, Schlievert PM, Bergdoll MS, Novick RP. 1983. The toxic shock syndrome exotoxin structural gene is not detectably transmitted by a prophage. Nature 305:709–712.

46. Novick RP. 1991. [27] Genetic systems in Staphylococci, p 587-636, Methods in Enzymology, vol 204. Academic Press.

47. Rosario-Cruz Z, Chahal HK, Mike LA, Skaar EP, Boyd JM. 2015. Bacillithiol has a role in Fe-S cluster biogenesis in *Staphylococcus aureus*. Mol Microbiol 98:218–42.

48. Mashruwala AA, Boyd JM. 2015. De Novo Assembly of Plasmids Using Yeast Recombinational Cloning. Methods Mol Biol 1373:33–41.

49. Joska TM, Mashruwala A, Boyd JM, Belden WJ. 2014. A universal cloning method based on yeast homologous recombination that is simple, efficient, and versatile. J Microbiol Methods 100:46–51.

50. Mashruwala AA, Bhatt S, Poudel S, Boyd ES, Boyd JM. 2016. The DUF59 Containing Protein SufT Is Involved in the Maturation of Iron-Sulfur (FeS) Proteins during Conditions of High FeS Cofactor Demand in *Staphylococcus aureus*. PLoS Genet 12:e1006233.

51. Olson BJ, Markwell J. 2007. Assays for determination of protein concentration. Curr Protoc Protein Sci Chapter 3:Unit 3 4.

52. Carroll RK, Weiss A, Shaw LN. 2016. RNA-Sequencing of Staphylococcus aureus Messenger RNA. Methods Mol Biol 1373:131–41.

53. Carroll RK, Weiss A, Broach WH, Wiemels RE, Mogen AB, Rice KC, Shaw LN. 2016. Genome-wide Annotation, Identification, and Global Transcriptomic Analysis of Regulatory or Small RNA Gene Expression in Staphylococcus aureus. mBio 7:e01990–15.

54. Tatusov RL, Galperin MY, Natale DA, Koonin EV. 2000. The COG database: a tool for genome-scale analysis of protein functions and evolution. Nucleic Acids Res 28:33–6.

55. Bastock RA, Marino EC, Wiemels RE, Holzschu DL, Keogh RA, Zapf RL, Murphy ER, Carroll RK. 2021. Staphylococcus aureus Responds to Physiologically Relevant Temperature Changes by Altering Its Global Transcript and Protein Profile. mSphere 6.

56. Schmittgen TD, Livak KJ. 2008. Analyzing real-time PCR data by the comparative C(T) method. Nat Protoc 3:1101–8.

57. Yang JH, Wright SN, Hamblin M, McCloskey D, Alcantar MA, Schrubbers L, Lopatkin AJ, Satish S, Nili A, Palsson BO, Walker GC, Collins JJ. 2019. A White-Box Machine Learning Approach for Revealing Antibiotic Mechanisms of Action. Cell 177:1649–1661 e9.

58. Shlomi T, Cabili MN, Herrgard MJ, Palsson BO, Ruppin E. 2008. Network-based prediction of human tissue-specific metabolism. Nat Biotechnol 26:1003–10.

59. Zur H, Ruppin E, Shlomi T. 2010. iMAT: an integrative metabolic analysis tool. Bioinformatics 26:3140–2.

60. Ebrahim A, Lerman JA, Palsson BO, Hyduke DR. 2013. COBRApy: COnstraints-Based Reconstruction and Analysis for Python. BMC Syst Biol 7:74.

61. Megchelenbrink W, Huynen M, Marchiori E. 2014. optGpSampler: an improved tool for uniformly sampling the solution-space of genome-scale metabolic networks. PLoS One 9:e86587.

62. Patel JS, Norambuena J, Al-Tameemi H, Ahn YM, Perryman AL, Wang X, Daher SS, Occi J, Russo R, Park S, Zimmerman M, Ho HP, Perlin DS, Dartois V, Ekins S, Kumar P, Connell N, Boyd JM, Freundlich JS. 2021. Bayesian Modeling and Intrabacterial Drug Metabolism Applied to Drug-Resistant Staphylococcus aureus. ACS Infect Dis 7:2508–2521.

63. Melamud E, Vastag L, Rabinowitz JD. 2010. Metabolomic analysis and visualization engine for LC-MS data. Anal Chem 82:9818–26.

64. Schwyn B, Neilands JB. 1987. Universal chemical assay for the detection and determination of siderophores. Anal Biochem 160:47–56.

65. Arora NK, Verma M. 2017. Modified microplate method for rapid and efficient estimation of siderophore produced by bacteria. 3 Biotech 7:381.

66. Buchfink B, Reuter K, Drost HG. 2021. Sensitive protein alignments at tree-of-life scale using DIAMOND. Nat Methods 18:366–368.

67. Katoh K, Kuma K, Toh H, Miyata T. 2005. MAFFT version 5: improvement in accuracy of multiple sequence alignment. Nucleic Acids Res 33:511–8.

68. Lefort V, Desper R, Gascuel O. 2015. FastME 2.0: A Comprehensive, Accurate, and Fast Distance-Based Phylogeny Inference Program. Mol Biol Evol 32:2798–800.

69. Lemoine F, Gascuel O. 2021. Gotree/Goalign: toolkit and Go API to facilitate the development of phylogenetic workflows. NAR Genom Bioinform 3:lqab075.

70. Letunic I, Bork P. 2021. Interactive Tree Of Life (iTOL) v5: an online tool for phylogenetic tree display and annotation. Nucleic Acids Res 49:W293–W296.

71. Galili T. 2015. dendextend: an R package for visualizing, adjusting and comparing trees of hierarchical clustering. Bioinformatics 31:3718–20.

72. Moriya Y, Itoh M, Okuda S, Yoshizawa AC, Kanehisa M. 2007. KAAS: an automatic genome annotation and pathway reconstruction server. Nucleic Acids Res 35:W182–5.

73. Jumper J, Evans R, Pritzel A, Green T, Figurnov M, Ronneberger O, Tunyasuvunakool K, Bates R, Zidek A, Potapenko A, Bridgland A, Meyer C, Kohl SAA, Ballard AJ, Cowie A, Romera-Paredes B, Nikolov S, Jain R, Adler J, Back T, Petersen S, Reiman D, Clancy E, Zielinski M, Steinegger M, Pacholska M, Berghammer T, Bodenstein S, Silver D, Vinyals O, Senior AW, Kavukcuoglu K, Kohli P, Hassabis D. 2021. Highly accurate protein structure prediction with AlphaFold. Nature 596:583–589.

74. Greco TM, Miteva Y, Conlon FL, Cristea IM. 2012. Complementary proteomic analysis of protein complexes. Methods Mol Biol 917:391–407.

75. Tsai YC, Greco TM, Boonmee A, Miteva Y, Cristea IM. 2012. Functional proteomics establishes the interaction of SIRT7 with chromatin remodeling complexes and expands its role in regulation of RNA polymerase I transcription. Mol Cell Proteomics 11:M111.015156.

76. Wiśniewski JR, Zougman A, Nagaraj N, Mann M. 2009. Universal sample preparation method for proteome analysis. Nat Methods 6:359–62.

77. Manza LL, Stamer SL, Ham AJ, Codreanu SG, Liebler DC. 2005. Sample preparation and digestion for proteomic analyses using spin filters. Proteomics 5:1742–5.

78. Erde J, Loo RR, Loo JA. 2014. Enhanced FASP (eFASP) to increase proteome coverage and sample recovery for quantitative proteomic experiments. J Proteome Res 13:1885–95.

79. Carabetta VJ, Greco TM, Cristea IM, Dubnau D. 2019. YfmK is an Ne-lysine acetyltransferase that directly acetylates the histone-like protein HBsu in *Bacillus subtilis*. Proceedings of the National Academy of Sciences 116:3752–3757.

80. Masuda T, Tomita M, Ishihama Y. 2008. Phase transfer surfactant-aided trypsin digestion for membrane proteome analysis. Journal of Proteome Research 7:731–740.

81. Kulak NA, Pichler G, Paron I, Nagaraj N, Mann M. 2014. Minimal, encapsulated proteomic-sample processing applied to copy-number estimation in eukaryotic cells. Nature Methods 11:319–324.

82. Ashford P, Hernandez A, Greco TM, Buch A, Sodeik B, Cristea IM, Grünewald K, Shepherd A, Topf M. 2016. HVint: A strategy for identifying novel protein-protein interactions in Herpes Simplex Virus Type 1. Mol Cell Proteomics 15:2939–53.

83. Grossmann J, Roschitzki B, Panse C, Fortes C, Barkow-Oesterreicher S, Rutishauser D, Schlapbach R. 2010. Implementation and evaluation of relative and absolute quantification in shotgun proteomics with label-free methods. J Proteomics 73:1740–6.

84. Kuzmič P. 1996. Program DYNAFIT for the Analysis of Enzyme Kinetic Data: Application to HIV Proteinase. Analytical Biochemistry 237:260–273.

85. George GN, George SJ, Pickering IJ. 2001. EXAFSPAK, http://www-ssrl.slac.stanford.edu/~george/exafspak/exafs.htm, Menlo Park, CA.

86. Ankudinov AL, Rehr JJ. 1997. Relativistic Spin-dependent X-ray Absorption Theory. Phys Rev B 56:1712.

87. Cook JD, Kondapalli KC, Rawat S, Childs WC, Murugesan Y, Dancis A, Stemmler TL. 2010. Molecular details of the yeast frataxin-Isu1 interaction during mitochondrial Fe-S cluster assembly. Biochemistry 49:8756–65.

88. Riggs-Gelasco PJ, Stemmler TL, Penner-Hahn JE. 1995. XAFS of Dinuclear Metal Sites in Proteins and Model Compounds. Coord Chem Review 114.

89. Fey PD, Endres JL, Yajjala VK, Widhelm TJ, Boissy RJ, Bose JL, Bayles KW. 2013. A genetic resource for rapid and comprehensive phenotype screening of nonessential *Staphylococcus aureus* genes. MBio 4:e00537–12.

90. Mashruwala AA, Roberts CA, Bhatt S, May KL, Carroll RK, Shaw LN, Boyd JM. 2016. *Staphylococcus aureus* SufT: An essential iron-sulfur cluster assembly factor in cells experiencing a high-demand for lipoic acid. Mol Microbiol doi:10.1111/mmi.13539.

91. Mashruwala AA, Van De Guchte A, Boyd JM. 2017. Impaired respiration elicits SrrAB-dependent programmed cell lysis and biofilm formation in *Staphylococcus aureus*. eLife 6.

92. Albano M, Hahn J, Dubnau D. 1987. Expression of competence genes in Bacillus subtilis. J Bacteriol 169:3110–7.

93. Forsyth RA, Haselbeck RJ, Ohlsen KL, Yamamoto RT, Xu H, Trawick JD, Wall D, Wang L, Brown-Driver V, Froelich JM, C KG, King P, McCarthy M, Malone C, Misiner B, Robbins D, Tan Z, Zhu Zy ZY, Carr G, Mosca DA, Zamudio C, Foulkes JG, Zyskind JW. 2002. A genome-wide strategy for the identification of essential genes in *Staphylococcus aureus*. Mol Microbiol 43:1387–400.

94. Pang YY, Schwartz J, Thoendel M, Ackermann LW, Horswill AR, Nauseef WM. agr-Dependent interactions of Staphylococcus aureus USA300 with human polymorphonuclear neutrophils. J Innate Immun 2:546–59.

95. Bose JL, Fey PD, Bayles KW. 2013. Genetic tools to enhance the study of gene function and regulation in *Staphylococcus aureus*. Appl Environ Microbiol 79:2218–24.

96. Kramer N, Hahn J, Dubnau D. 2007. Multiple interactions among the competence proteins of *Bacillus subtilis*. Mol Microbiol 65:454–64.

97. Bubeck Wardenburg J, Williams WA, Missiakas D. 2006. Host defenses against Staphylococcus aureus infection require recognition of bacterial lipoproteins. Proc Natl Acad Sci U S A 103:13831–6.

